# Reducing spatial heterogeneity in coverage improves the effectiveness of dog vaccination campaigns against rabies

**DOI:** 10.1101/2024.10.03.616420

**Authors:** Elaine A Ferguson, Ahmed Lugelo, Anna Czupryna, Danni Anderson, Felix Lankester, Lwitiko Sikana, Jonathan Dushoff, Katie Hampson

## Abstract

Vaccination programs are the mainstay of control for many infectious diseases. Heterogeneous coverage is hypothesised to reduce vaccination programme effectiveness, but this impact has not been quantified in real systems. We address this gap using fine-scale data from two decades of rabies contact tracing and dog vaccination campaigns in Serengeti district, Tanzania. We also aimed to identify drivers of continued circulation of rabies in the district despite annual vaccination campaigns. Using generalised linear mixed models, we find that current focal (village-level) dog rabies incidence decreases with increasing recent focal vaccination coverage. However, current focal incidence depends most on recent incidence, both focally and in the wider district, consistent with high population connectivity. Removing the masking effects of prior non-focal incidence shows that, for the same average prior non-focal (wider-district) vaccination coverage, increased heterogeneity in coverage among the non-focal villages leads to increased focal incidence. These effects led to outbreaks following years when vaccination campaigns missed many villages, whereas when heterogeneity in coverage was reduced, incidence declined to low levels (<0.4 cases/1,000 dogs annually and no human deaths) and short vaccination lapses thereafter did not lead to resurgence. Through transmission-tree reconstruction, we inferred frequent incursions into the district each year (mean of 7). Inferred incursions substantially increased as a percentage of all cases in recent years, reaching 50% in 2022, suggesting regional connectivity is driving residual transmission. Overall, we empirically demonstrate how population connectivity and spatial heterogeneity in vaccination can impact disease outcomes, highlighting the importance of fine-scale monitoring in managing vaccination programs.

## Introduction

Vaccination of domestic dogs to interrupt rabies transmission is a quintessential One Health intervention that mitigates risks of human infection from this fatal zoonotic disease [1,2]. While the impacts of vaccinating human populations against childhood diseases have been studied extensively, vaccination of animal populations has not been examined in the same detail, and impacts may differ for a number of reasons. Firstly, demographic turnover and therefore waning of vaccine-induced immunity is likely to be faster in many animal populations than in humans [3–5]; secondly, with the exception of species that undertake long-range migrations [6], animal movement is often more local, perhaps curtailing the geographic spread of disease [7]; and thirdly, vaccinating animals is potentially more difficult, with logistical challenges resulting in lower and/or more heterogeneous coverage [8–11]. The implications of these differences are important to understand for rabies, which kills tens of thousands of people every year and causes billions of dollars of economic losses [12]. Moreover, understanding how vaccination coverage impacts transmission in animal populations could have far-reaching implications for a range of zoonoses and for diseases that threaten food security and endangered wildlife.

Rabies is a fatal viral disease of mammals, with dog-mediated rabies being the cause of >99% of human rabies deaths [13]. Post-exposure vaccines are effective in preventing infection onset, but are costly for both the health sector and bite victims, and lack of access to these emergency vaccines has tragic consequences [14–16]. However, rabies can be tackled at source through mass dog vaccination. When sufficient coverage is reached, dog vaccination can interrupt transmission and even eliminate rabies, with economic benefits from the reduced need for post-exposure vaccines [17–19]. Dog vaccination is therefore the cornerstone of the global strategic plan, ‘Zero by 30’, to end human deaths from dog-mediated rabies by 2030 [20].

Rabies control by mass dog vaccination is not a new concept. Dog vaccination was integral to the elimination of rabies from Japan in 1957, and is still maintained as a safeguard against reintroductions [21]. Mass dog vaccination was also central to reducing dog-mediated rabies by >95% between 1980 and 2010 in Latin America and the Caribbean [22], with Mexico becoming the first country validated by WHO as free from dog-mediated human rabies since the ‘Zero by 30’ initiative began [23]. Despite these huge strides in the Americas, mass dog vaccination has not been implemented at scale across Africa and Asia. Pilot vaccination projects have reduced local rabies incidence [24–27], dispelled doubts about the accessibility of free-roaming dogs for vaccination, elucidated their role as the reservoir population [28], and provided valuable lessons for improving vaccination campaign design [29–31]. However, piecemeal vaccination efforts are not often maintained or rolled out nationally due to low prioritisation of this neglected disease [32]. Lack of consistent, large-scale vaccination may have negative consequences for the long-term impacts of local efforts. Areas where rabies has been eliminated, but where dog vaccination has lapsed, are vulnerable to reintroductions, as observed in many settings [26,33,34]. However, relatively little is known about rates of incursions and the distances over which they can penetrate into control areas. Better understanding of incursion risks would inform, for example, how wide vaccinated buffer zones must be to prevent introductions from establishing. Further knowledge of how incursions, local dog movements, and vaccination efforts interact to drive disease dynamics would also provide valuable guidance to rabies-endemic countries as they develop strategic plans to reach the ‘Zero by 30’ goal.

Assessments of mass dog vaccination campaigns suggest that coverage can be highly heterogeneous [29,35,36], often failing to reach the recommended 70% target [25,37]. Reasons for these outcomes include lack of resources and insufficient attention to barriers affecting participation [9,10,35]. This is concerning, as simulation-based models indicate that patches of low coverage could jeopardise rabies control more widely [36,38]. However, the impacts of heterogeneous coverage have rarely, if ever, been quantified using incidence data from a real system, rabies or otherwise. Fine-scale heterogeneities in vaccination coverage may be masked by data aggregated to administrative scales. Such aggregation has been a feature of most empirical studies reporting impacts of rabies vaccination [25,26,39,40] and limits our understanding of how incidence is driven by influences at both local and wider scales. Fine-scale data could allow quantification of these impacts, revealing the consequences of coverage heterogeneities and the extent of epidemiological connectivity.

Here we examine dog rabies and human rabies exposures and deaths in the Serengeti district of northern Tanzania, where dog vaccination has been ongoing since 2003, along with contact tracing to track rabies transmission. It is unclear how rabies has been able to persist in Serengeti district despite control efforts, and in this study we explore evidence for three hypotheses: 1) Spatial heterogeneity in vaccination within the district allows rabies to persist in patches of lower coverage; 2) The mean vaccination coverage over the district has simply been insufficient; 3) Incursions of rabies from outside the district maintain transmission. Using contact tracing and vaccination campaign data at fine spatiotemporal scales to inform statistical models, we decipher the cross-scale drivers of viral circulation and draw conclusions on what is necessary to eliminate rabies from an animal reservoir in a connected landscape.

## Results

### Study area and dog population

Serengeti district consists of 88 villages, each of which shares borders with a median of 5 (range: 2-9) neighbouring villages. Village polygons ranged in area from 5.5-386.2km^2^, but numbers of cells on a 1km^2^ grid (Fig. 1A) with a non-zero population (from a georeferenced district-wide census of humans and dogs) ranged from 5 to just 85. The district is bordered both by other districts where rabies circulates endemically, and by Serengeti National Park where domestic dogs are not permitted (Fig. 1A).

**Figure 1.**
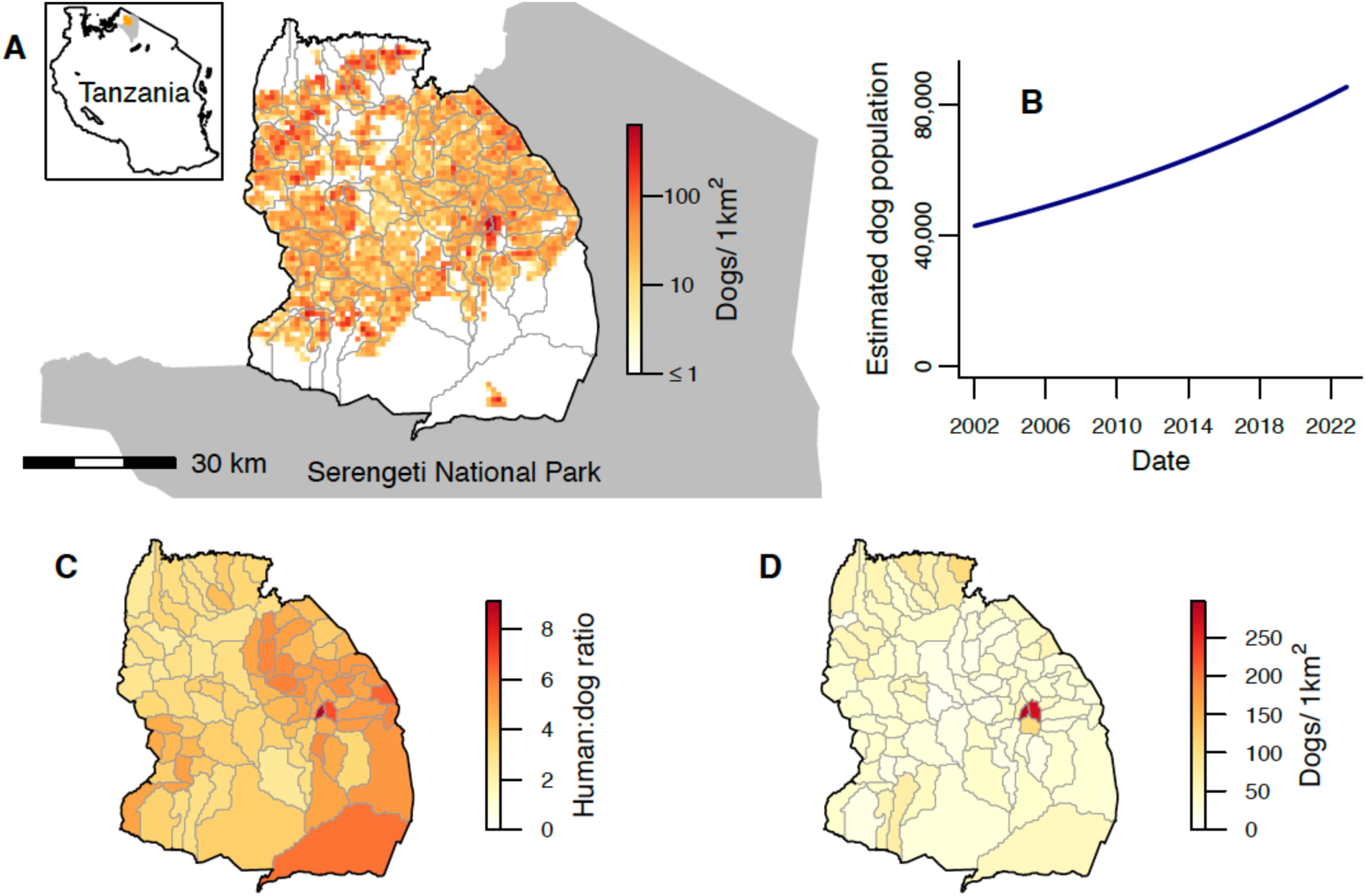
Study area and dog population. A) Serengeti District bordered by Serengeti National Park (grey shading). Grey lines show village borders within the district. The spatial distribution of the estimated dog population on a 1km^2^ grid at the end of 2022 is indicated by the colour scale (on a log scale). The inset indicates the location of Serengeti District (blue) and Serengeti National Park (grey) within Tanzania. B) Total dog population in Serengeti District from 2002-2022, estimated based on national human census data and human:dog ratios. C) Human:dog ratios for each village from a district-wide census of humans and dogs. D) Village-level dog density at the end of 2022, estimated by dividing the estimated village dog populations by the total number of 1km^2^ cells with non-zero population in each village.

Serengeti district’s human population grew from an estimated 175,017 to 345,587 between January 2002 and December 2022 based on national census data; an average growth rate of 3.2% per annum. Human:dog ratios (estimated from the georeferenced human and dog census) varied from 2.4-9.1 between villages (Fig. 1C) [41]. The district dog population, estimated from these ratios, approximately doubled over the study period, increasing from 42,931 to 85,672 (Fig. 1B). Estimated dog densities in December 2022 ranged from 0-625 dogs/km^2^ across a 1km^2^ grid (Fig. 1A, mean of 21 dogs/km^2^ or 37 dogs/km^2^ when considering only the 57% of cells with a non-zero dog population). Village-level dog densities, which ranged from 10-297 dogs/km^2^ in December 2022 (Fig. 1D), were calculated by dividing estimated village dog populations by the number of 1km^2^ cells with non-zero population in the village, to better reflect densities experienced by populations in villages with large unoccupied areas, particularly on the North and South borders (Fig. 1A).

### Heterogeneous vaccination coverage

Annual rabies vaccination campaigns were initiated as part of a research project in 2003 (Fig. 2A) in the district’s villages within 10 kilometres of the Serengeti National Park [42,43]. The percentage of villages that held campaigns in 2003, hereafter “campaign completeness”, was 48% (Fig. 2A,E, Table S1). In 2004, in response to a rabies outbreak [44], campaigns were expanded by the local government, resulting in a campaign completeness of 90% (Fig. 2A,D-E). Subsequently, campaigns continued in villages bordering the National Park, and less consistently across the rest of the district, depending on local government resources. Following a resurgence of rabies in 2010-2012, the local government committed to the goal of conducting campaigns annually in all villages throughout the district. Campaign completeness increased gradually, reaching ≥95% throughout 2015-2017. In 2018, vaccination in the northwest villages lapsed again due to logistical constraints (Fig. 2A,E). But, in 2019, for the first time, campaign completeness reached 100%. In 2021, vaccination did not take place in the southeastern villages for the first time since 2003, however as the 2020 campaigns in these villages happened late in the calendar year and the 2022 campaigns were early, the inter-campaign interval was only 16-18 months. The mean campaign completeness over all years from 2003-2022 was 77%.

**Figure 2.**
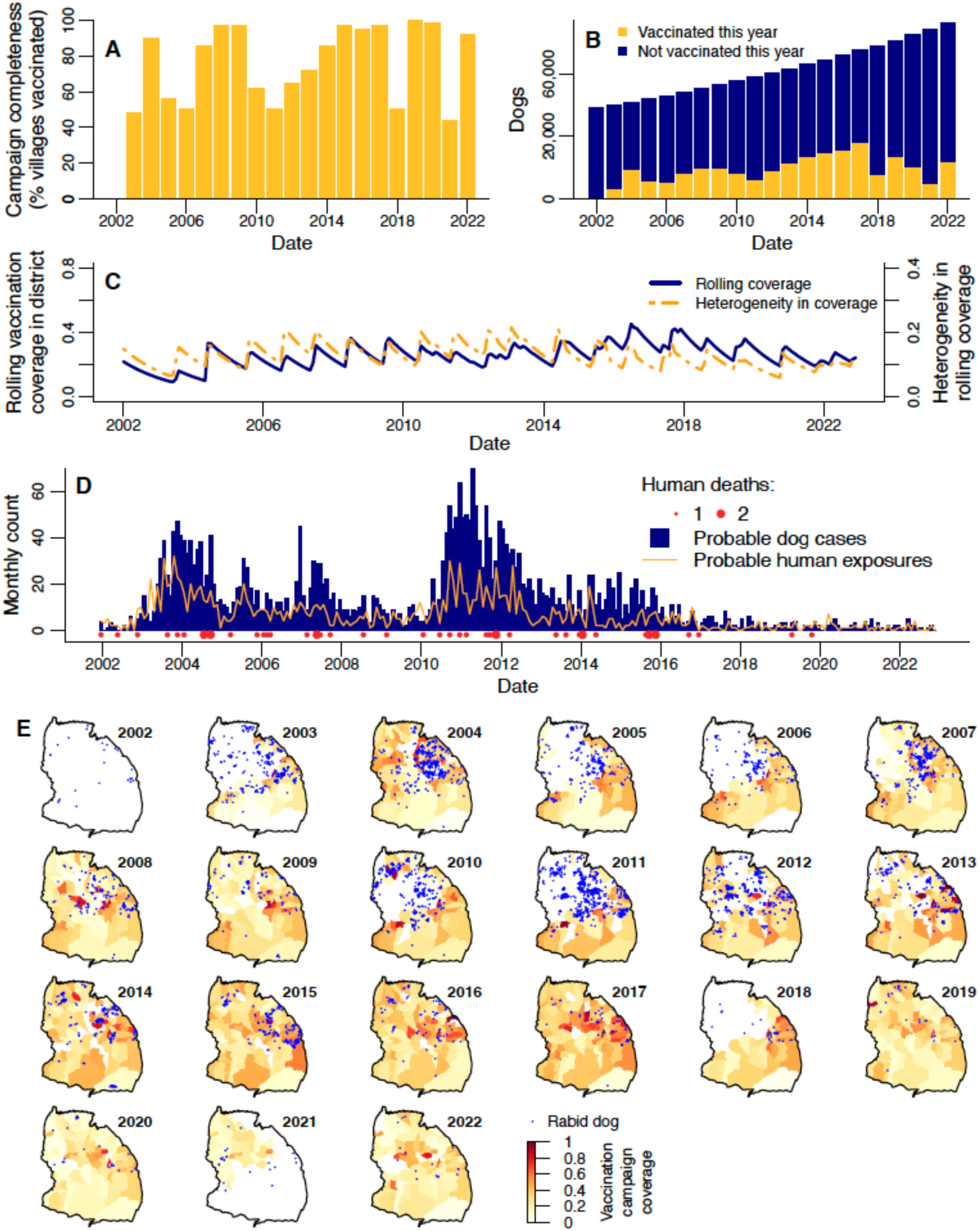
Dog vaccination, dog rabies cases and human rabies exposures and deaths in Serengeti district from 2002-2022. A) Campaign completeness (% villages in the district that held vaccinations) each year. B) Number of dogs that were and were not vaccinated in the district each year. C) Monthly district-level ‘rolling’ dog vaccination coverage (blue line), and monthly heterogeneity in rolling coverage (the population-weighted standard deviation of rolling coverage over all villages in the district; orange line). D) Monthly dog cases, human exposures, and human deaths in the district. E) Village-level ‘campaign’ vaccination coverage (colour scale) and locations of dog rabies cases (blue points) each year (see Fig. S1B for campaign coverage maps without overlaid cases).

Numbers of dogs vaccinated in campaigns each year ranged from 4,199 in 2003 to 26,419 in 2017 (Fig. 2B). Like campaign completeness, dogs vaccinated steadily increased each year following the local government commitment to annual campaigns. However, since the peak in 2017, numbers of dogs vaccinated each year have fallen again, despite the increasing dog population (Fig. 2B).

From 2003 until 2022, the percentage of dogs vaccinated during annual campaigns (hereafter “campaign coverage”) in the district ranged from 8-37%, with a mean of 22% (Fig. S1A, Table S1). Campaign coverage varied more across villages, from 0.3-98%, with a mean of 31% (Fig. 2E). Only 3% of village campaigns reached the recommended target coverage of 70%.

In each month in 2002-2022, we also estimated the proportion of dogs in the population that had been vaccinated at least once in their lives, accounting for vaccination campaign timing and dog population turnover. This “rolling coverage” estimate can exceed campaign coverage, since it includes vaccinated dogs that have survived from previous years but that may not have been re-vaccinated in the current year (we assume that previously vaccinated and unvaccinated dogs have an equal probability of being vaccinated in annual campaigns). Our estimate of district-level rolling coverage in January 2002 when the study started was 22% based on information from the 2000-2001 vaccination campaign preceding our study [25], with subsequent fluctuation between 9-45% (mean of 26%; Fig. 2B). For most of the study period, there was relatively little variation in the magnitude of district-level rolling coverage (Fig. 2B, Fig. S2A), but rolling coverage between villages was more variable (Video S1, Fig. S2). Specifically, coverage heterogeneity, calculated as the standard deviation of rolling coverage over all villages, weighted by village dog populations (Fig. 2B), was observed to be higher in 2007 and 2011-2013 (Fig. 2C, Fig. S2B). The peaks in heterogeneity followed multi-year periods (in 2002-2003, 2005-2006 and 2010-2012) where groups of villages in the northwest were not vaccinated. Single years where groups of villages were not vaccinated (in 2018 and 2021) did not lead to visually obvious drops in the yearly average rolling coverage in those areas (Fig. S2C), and this is reflected in relatively low levels of coverage heterogeneity from 2019-2022 (Fig. 2C, Fig. S2B).

### Rabies cases and exposures

A total of 3,362 probable dog rabies cases were identified from January 2002 to December 2022 via contact tracing. Identification of probable animal cases was based on the presence of clinical signs along with either: 1) disappearance or death of the animal within 10 days, or 2) the animal being killed and either of unknown origin or known to have previously been bitten [44]. Annually, probable case numbers varied from just 16 (0.19 cases/1,000 dogs) in 2022 up to 547 (9.4 cases/1,000 dogs) in 2011, while monthly cases ranged from 0 to 70 (Fig. 2D), never exceeding 17 in any village (Video S1). Dog cases occurred primarily in the central and northern villages, and were less frequent in the south (Fig. 2E). At least one case was recorded in every village, except for two neighbouring villages in the west of the district. Dog rabies incidence was relatively low in 2002 but increased over 2003, leading to an outbreak that was largely controlled by the district-wide vaccination campaign in 2004 (Fig. 2D-E). Further outbreaks (which we define as surges with cases exceeding 4/1,000 dogs/year, since this captures the most prominent peaks observed in the district time series) occurred in 2005 and 2007, during or following years without district-wide vaccination and culminated in the largest outbreak from 2010-2012. Cases steadily declined from 2012 and remained at low levels of <30 per year (<0.4 cases/1,000 dogs) from 2018 onwards despite lapses in vaccination campaigns in 2018 and 2021 (Fig. 2).

Probable human rabies exposures (individuals identified through contact tracing – via bite patient records from health facilities and subsequent interviews with patients/dog-owners – who received a bite/scratch from a probable rabies case) roughly tracked dog rabies, ranging from 0 to 32 exposures per month and peaking in late 2003 (Fig. 2D). Annual exposures ranged from 204 in 2003 (111 exposures/100,000 people) to just 11 in 2021 and 2022 (3 exposures/100,000 people), having declined and remained low (≤21 per year) since 2017.

Individual rabid dogs exposed between 0-18 people (mean=0.39, Fig. S3), with 72% not exposing anyone, and 21% exposing just one person. Of all human exposures, 87.5% were due to domestic dogs. Exposures from other species followed a similar temporal pattern to those by dogs (Fig. S4). Of non-dog-mediated human exposures where the biting species was known, 47.9% were by domestic cats, 18.9% by jackals, 11.6% by livestock, 2.1% by humans, and the remaining 19.5% by various other wildlife species. In comparison, 84.6% of rabid animals identified by contact tracing were domestic dogs, with the remainder being composed of 60.5% livestock, 13.3% domestic cats, 12.0% jackals and 14.1% various other wildlife species.

Overall, 48 human rabies deaths were identified (Fig. 2D, Video S1), with a peak of seven in 2011 (annual incidence of 3 deaths/100,000). Deaths were recorded every year until 2018. Two deaths were recorded again in 2019 despite low numbers of exposures and dog cases, but there were no deaths thereafter. Of the 48 deaths, none completed a full course of post-exposure vaccinations: 3 individuals received one vaccination, 4 received two, and the remaining 41 received none.

### Drivers of disease incidence

To understand the drivers of these infection patterns we used negative binomial generalised linear mixed models (GLMMs). We modelled current focal incidence (cases/dog, i.e. cases in a village, with the logged village dog population as an offset) in response to prior rolling vaccination coverage and prior incidence (cases/dog), with a random effect of village. Both prior rolling coverage and prior incidence were included as averages over the previous two months, since >90% of rabies incubation periods recorded during contact tracing were shorter than two months (Fig. 2B). Prior rolling coverage and incidence were also considered at three spatial scales: 1) the focal village; 2) bordering villages; and 3) non-bordering villages in the district. We fitted models where prior rolling coverages in bordering and non-bordering villages were incorporated as either arithmetic means of rolling coverage or power means of susceptibility (i.e. 1 - rolling coverage) over those villages. Taking the power mean of susceptibility gives a measure of ‘effective susceptibility’ that is adjusted up or down from a simple arithmetic mean based on the degree of heterogeneity in susceptibility among the villages and the value of the parameter *p* (equation 13). If *p*>1, the power mean is biased towards higher susceptibility villages and rises above the arithmetic mean; i.e. for the same total number of unvaccinated dogs, the effective susceptibility is higher when those dogs are distributed unevenly. At *p*<1, heterogeneity reduces effective susceptibility, and at *p*=1, we revert to an arithmetic mean model. Since prior incidence at each spatial scale is expected to be partially determined by coverage/susceptibility at that scale (leading to correlation between these variables), including both variables in the model is expected to lead to partial masking of the vaccination effect. To better quantify the full independent impact of vaccination, we therefore fitted versions of the models both with and without the effects of prior incidence at each spatial scale (as specified in Table 1).

**Table 1:**
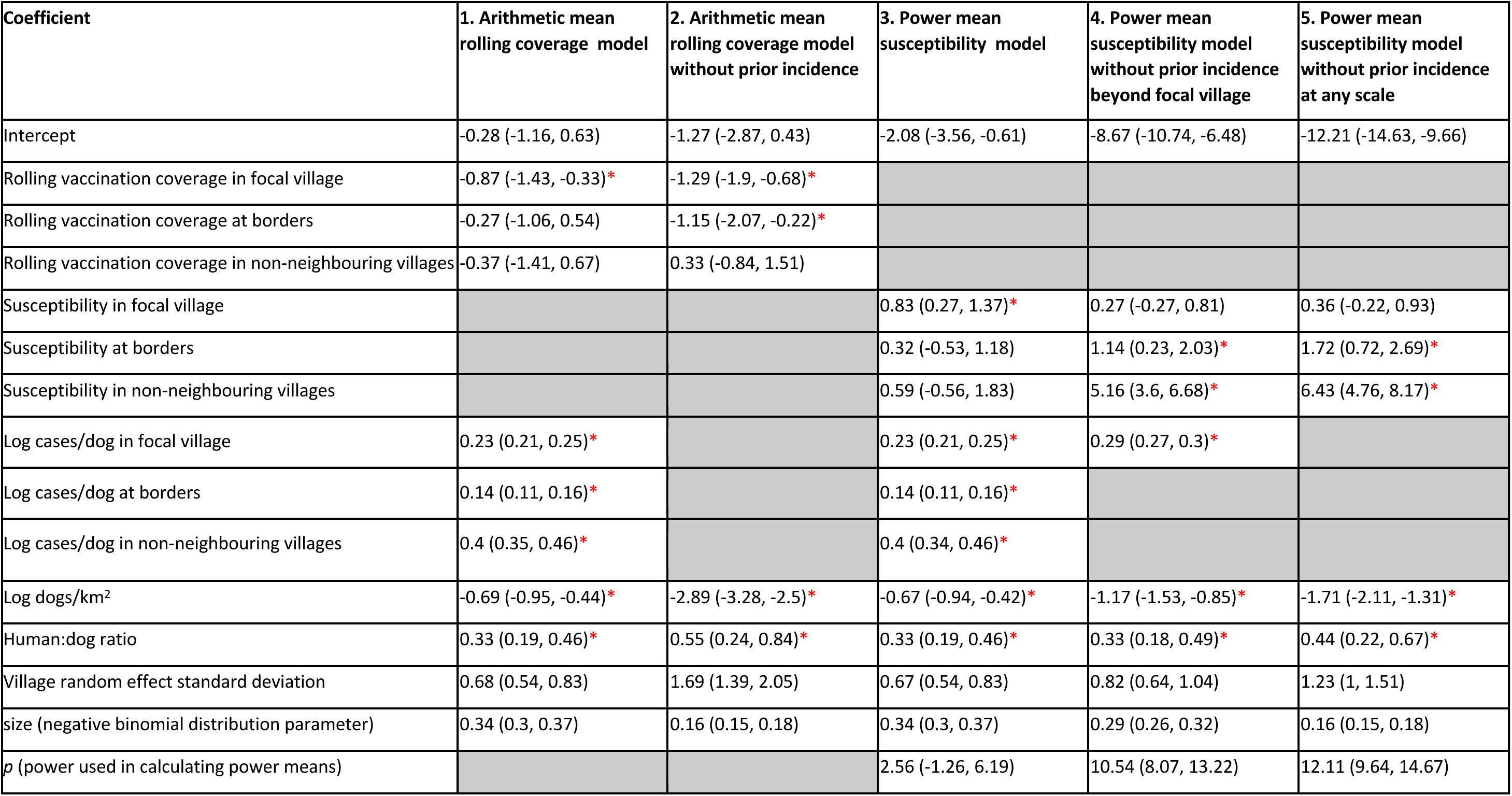
Coefficients for monthly village-level GLMMs for cases/dog in a village in the current month. 95% credible intervals (CrIs) in brackets. Coefficients for fixed effects where the 95% CrI does not include zero are marked *. Model 1 is presented in Fig. 3, model 2 in Fig. S3, and models 3 and 4 in Fig. 4. Vaccination, susceptibility and cases/dog variables are all averages over the prior two months.

Current focal incidence decreased with increasing prior focal rolling coverage (Fig. 3A, Table 1, Fig. S5A). In the arithmetic mean rolling coverage model with prior incidence, a 10% coverage increase in the focal village was associated with a focal incidence decrease of 8.3% (95% credible interval (CrI): 3.2-13.3%). In the arithmetic mean rolling coverage model without prior incidence, the estimated impact of a 10% increase in prior focal coverage was larger, causing a 13.4% (95% CrI: 8.3-18.2%) reduction in focal incidence. An impact of prior arithmetic mean rolling coverage in bordering villages was only evident in the model without prior incidence, while an effect of rolling coverage in non-bordering villages was not detected in either arithmetic mean rolling coverage model (Table 1, Fig. 3A, Fig. S5A).

**Figure 3:**
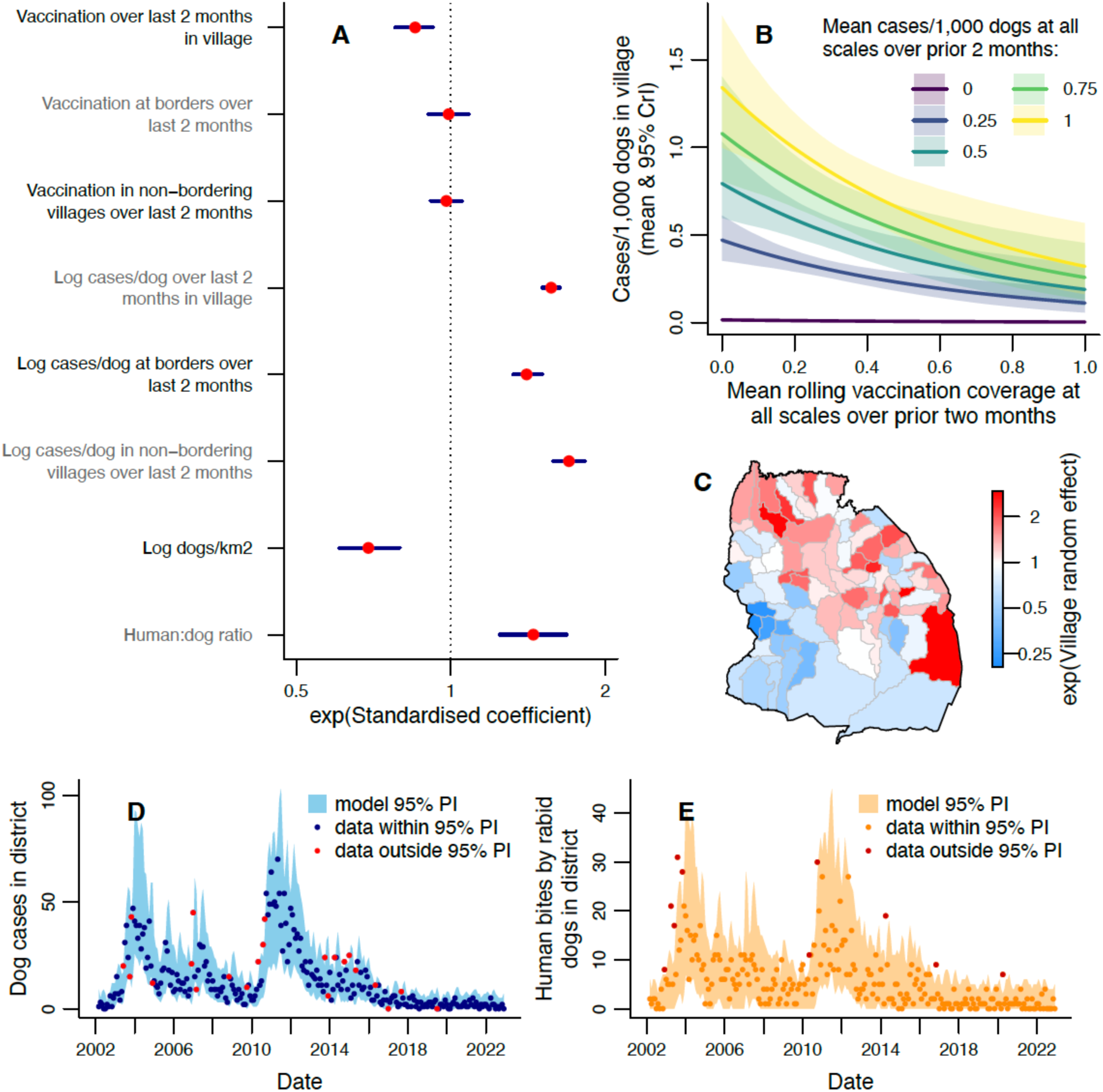
Modelling monthly dog rabies cases in villages. A) Exponentiated standardised values of the coefficients estimated for each explanatory variable, with 95% CrIs. See Table 1 for tabulated parameter values. B) Expected cases/1,000 dogs (number of dog cases normalised by dog population) in a village this month for different mean rolling vaccination coverages and mean cases/dog in the prior 2 months. Prior cases/dog values were chosen to represent the range observed at district level and shaded areas show 95% credible intervals (CrIs), with predictions obtained using average values of unspecified explanatory variables. C) Exponentiated random effect values for each village. D) Comparison of observed monthly dog cases (points) with the 95% prediction interval from the fitted model. E) Human rabies exposures each month (points), with 95% prediction interval from the dog case predictions in D and the fitted distribution of exposures per dog (Fig. S11). Data points in red (D, E) fall outside the model 95% prediction interval (PI).

We find that incidence increases with power mean susceptibility in bordering and non-bordering villages only when prior incidence at those scales is not included (Table 1, Fig. 4A). In those models we estimate *p* to be substantially larger than one (Table 1, Fig. 4B) and beyond what we *a priori* expected to be the feasible range (Fig. S6). The increase in effective susceptibility at the district level due to heterogeneity (Fig. 4C, Fig. S8) follows a similar pattern to the standard deviation in village coverage (Fig. 2B, Fig. S2B), having peaks in 2007 and 2013, and being lower from 2019.

**Figure 4:**
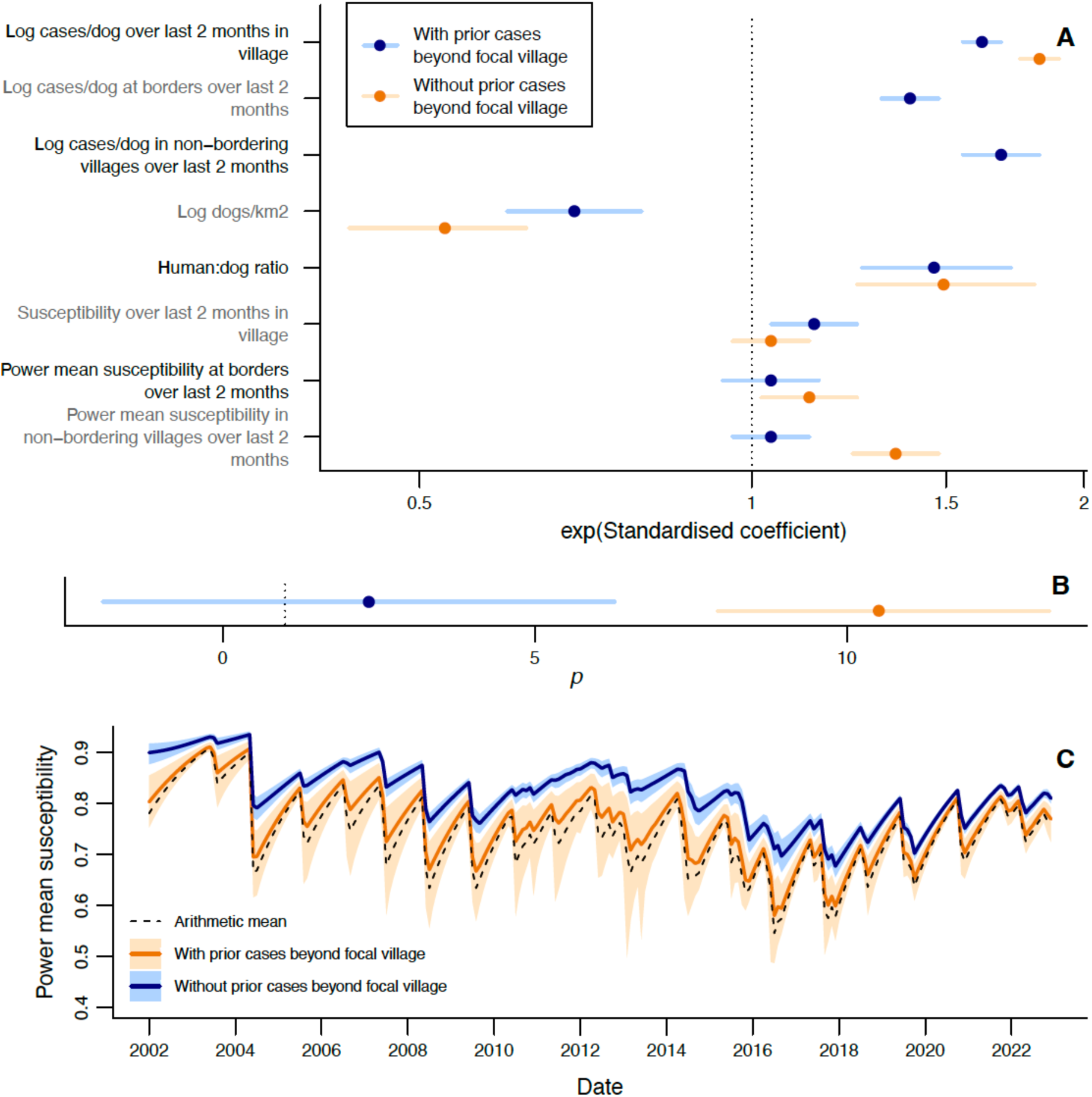
Using power mean susceptibility to model impacts of heterogeneity in vaccination on rabies incidence at the village level. A) Exponentiated standardised estimated coefficients for explanatory variables, with 95% CrIs. B) Estimated power *p* used to calculate power mean susceptibilities. A-B show estimates from models with (blue) and without (orange) effects of prior incidence beyond the focal village. See Table 1 for tabulated parameter values. C) Monthly arithmetic mean susceptibility over all villages in the district, compared with power mean susceptibility (mean and 95% CrI) from fitted values of *p* from models with and without prior incidence beyond the focal village.

The power mean susceptibility model without prior incidence beyond the focal village predicts, for example, that if half the bordering and non-bordering villages had prior coverage of 30%, with the other half having 50%, then the focal village incidence would be 1.4 (95% CrI: 1.3-1.6) times greater than if prior coverage had been homogeneous at 40%. If we further increase heterogeneity by assuming half the villages had prior 20% coverage and the other half 60%, then focal incidence is expected to be 2.6 (95% CrI: 1.9-3.4) times the homogeneous scenario.

Increased prior incidence across all three scales was associated with increased focal incidence (Fig. 3A-B, Fig. 5A) and had a greater impact than prior coverage or susceptibility (Fig. 3A, Fig. 5A), based on the standardised coefficients (found by transforming all explanatory variables to have a variance of one). Logging the prior incidence explanatory variables improved the fit based on the widely applicable information criterion (WAIC), a Bayesian model selection statistic [45]. Prior incidence in bordering villages had less impact on current focal incidence than prior incidence in non-bordering villages (Fig. 3A, Fig. 5A). In the arithmetic mean rolling coverage model, doubling mean cases/dog in the focal village over the prior two months led to a 17.3% (95% CrI: 15.7-18.9%) increase in cases/dog, while doubling at all three scales gave a 70.5% (95% CrI: 64.3-76.8%) increase.

**Figure 5.**
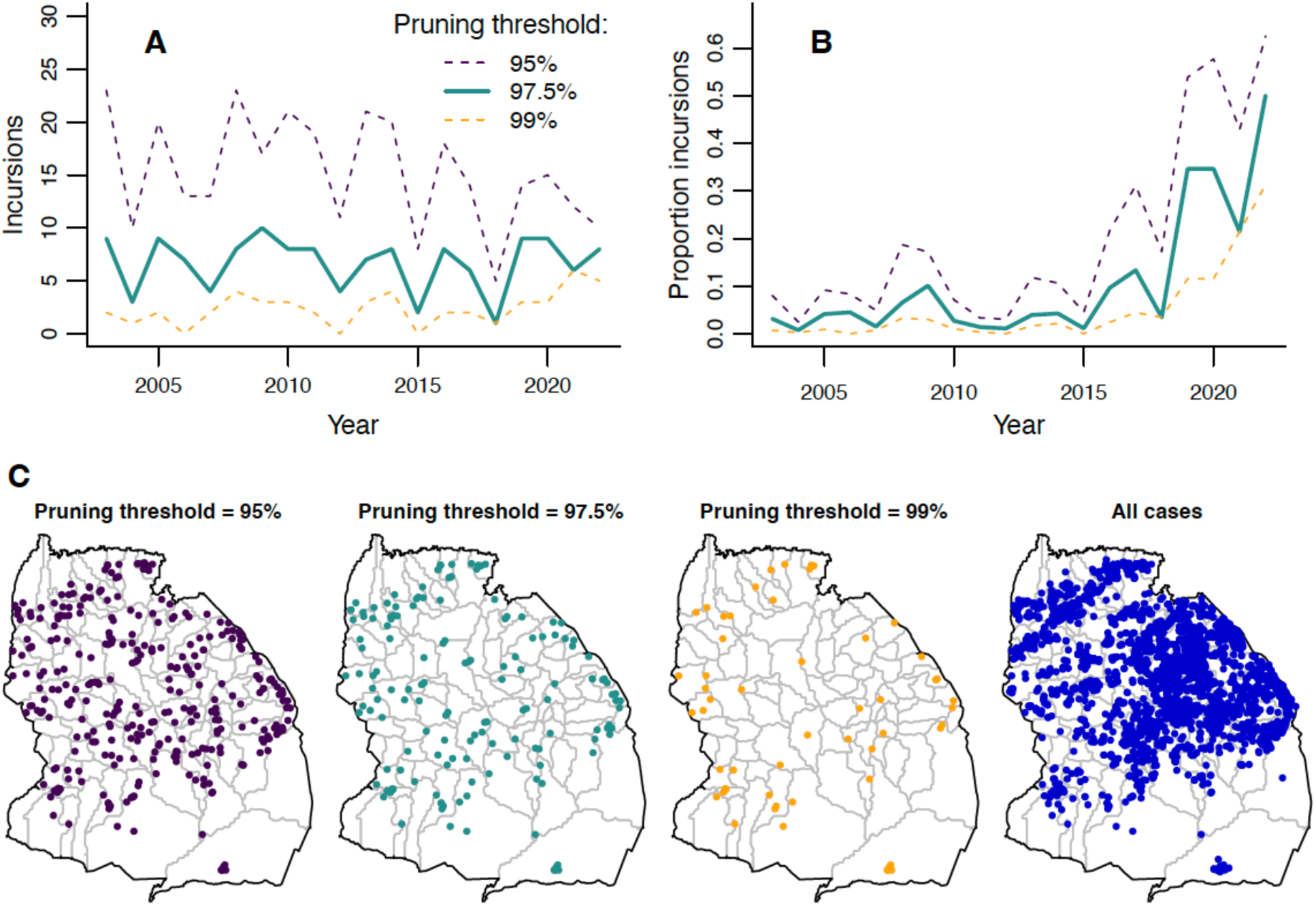
Estimated incursions in Serengeti District over time and space. A) Annual numbers of incursions inferred from transmission trees with three different pruning thresholds. B) Annual proportion of total rabies cases in carnivores that are inferred to be incursions. C) Locations of inferred incursions under the three pruning thresholds. Locations of all carnivore cases are shown for comparison.

An increase in focal incidence was associated with decreased dog density and increased human:dog ratio in the focal village (Fig. 3A, Fig. 4A). Random effects indicate that villages in the northeast of the district had higher than average incidence (Fig. 3C).

We modelled the remaining three combinations of village/district and monthly/annual scales. Models at district level consistently failed to detect impacts of prior vaccination, even on removal of prior incidence (Tables S2, S4, Figs S9-10, S12). Village-level annual models consistently detected effects of focal prior rolling coverage, but were inconsistent in detecting non-focal vaccination impacts (Table S3, Fig. S11). Models at both village and district level without prior incidence had a poorer fit to the data than the equivalent models with prior incidence (compare Fig. 3D to Fig. S5D and Fig. S7C to S7D).

A negative binomial distribution (mean=0.39 (±0.013, standard error); size=0.72 (±0.066)) fitted to the data on human exposures per rabid dog (Fig. S3) was used to simulate exposures from GLMM-based simulations of rabid dogs (Fig. 3D). This provided a good match to observed exposures (Fig. 3E).

By reconstructing transmission trees, we identified probable incursions into the district as cases without a plausible parent case within the 97.5th percentile of the distance kernel and serial interval distributions (Fig. S13, Table S5), i.e. the 97.5% pruning threshold. Numbers of incursions remained relatively constant from 2003-2022 (Fig. 5A, mean of 7 annual incursions). In contrast, the *proportion* of cases identified as incursions sharply increased (Fig. 5B), from 3% pre-2018 to 26% post-2018, peaking at 50% in 2022. Incursion locations were more focused along the district’s edges than cases in general (Fig. 5C). Adjusting the pruning threshold to 95% or 99% impacted the number of inferred incursions, but led to qualitatively similar temporal and spatial patterns (Fig. 5).

## Discussion

We identify important drivers of rabies transmission from twenty years of fine-scale data on dog vaccination and from contact tracing. Under endemic rabies circulation, we find that outbreaks occur following multi-year periods when clusters of villages remain unvaccinated. However, when district-wide vaccination was made routine (spurred by the largest outbreak in 2010-2012) and spatial heterogeneity in coverage reduced, incidence declined to low levels (<0.4 cases per 1,000 dogs annually), attributable largely to short chains of transmission following incursions into the district. Prior rabies incidence focally (at the scale of the focal village) and across the district was the main driver of current focal incidence, but was modulated by and correlated with prior focal vaccination coverage. When the masking effect of non-focal rabies incidence was removed, we also identified impacts of non-focal vaccination coverage and heterogeneity in this coverage. The role of prior incidence was highlighted in later years after vaccination had largely interrupted endemic transmission, when rabies did not re-surge despite late and incomplete vaccination campaigns. Reducing dog rabies cases had dramatic public health benefits, with corresponding reductions in human rabies exposures and deaths. However, frequent incursions suggest that benefits may be short-lived if dog vaccination lapses for extended periods.

Our analysis reveals how fine-scale variation in vaccination coverage drives rabies dynamics, with models aggregating coverage to district-level failing to show impacts on rabies incidence, demonstrating the limitations of coarsely aggregated data. Our empirical findings support simulation-based work arguing that gaps in coverage are detrimental for rabies control [36,38]. Pockets of low coverage are recognised as a driver of measles outbreaks in humans [46–49], but remain underexplored for other diseases, despite examples of vaccination heterogeneity in systems from cholera [50] to COVID-19 [51]. The impact of this heterogeneity on disease incidence has rarely, if ever, been quantified in real systems, likely due to a lack of fine-scale data. Our work addresses this gap, providing evidence that the same level of vaccination coverage can have substantially different impacts, based on its spatial distribution. Future studies may be aided by models that predict fine-scale vaccination coverage from coarser available data, e.g. sparse household surveys [52] or aggregated areal data [49]; though while these models may work well for routine childhood vaccinations, their transferability to the stochasticity arising from single-day village-level campaigns employed for dog rabies is unknown. The potential impact of clustered vs dispersed low-coverage villages also requires exploration. Nonetheless, striking 78.0-85.5% declines in district-level incidence were associated with a 35% increase in prior district-level coverage during large-scale rabies vaccination across 13 contiguous districts in south-east Tanzania, without accounting for heterogeneity [39]. These contrasting results were likely due to our focus on one district without wider-scale vaccination to reduce spread between districts. Our focus on a single, possibly idiosyncratic, time series may also be why the posterior for the parameter *p* that governs the heterogeneity effect lay outside the region that we *a priori* believed feasible; data from other areas could help refine estimates in future.

We find that vaccination controlled rabies despite not reaching the recommended 70% coverage target [25,37]. Some dog vaccination campaigns have faced similar difficulties in achieving high coverage [8,53], while others (even in Serengeti) have reported more success [25,54–56]. The low coverages we report may reflect our estimation methods [41]. We used vaccination records, with the dog population denominator derived from censuses and human:dog ratios. Methods like post-vaccination transects or household surveys may overestimate coverage if they rely on owner recall or do not cover unvaccinated sub-populations, resulting in bias towards more accessible areas and more visible (often adult) dogs [41,57]. Assuming constant human:dog ratios and a simple exponential human growth model in each village may have reduced the accuracy of our coverage estimates, while the relatively dense dog population and low human:dog ratio (versus [58] and [59]) may have contributed to the low coverage attained. Nonetheless, district coverage only fell below 20% – the critical threshold estimated to push the reproductive number below one [44] – on <5% of months post-2007 (Fig. 2C). So, while higher coverage would have achieved better (and faster) results, rabies was controlled regardless. Similarly, modelling suggests vaccination coverages of ≤40% could eliminate rabies [38] and protect against catastrophic declines in endangered canids [60]. We therefore conclude that, while high coverage should be the target for rapidly controlling disease, policymakers should not be deterred from introducing dog vaccination when achieving 70% coverage is challenging; if persistent spatial gaps are minimal, sub-optimal coverage still has major benefits due to rabies’ low transmissibility [44] relative to human diseases that require higher levels of vaccination [61]. However, fast demographic rates still necessitate many dogs be vaccinated annually to maintain coverage, in comparison to childhood vaccination programmes where higher coverage is reached through targeting a narrower demographic [62].

Impacts of prior rabies incidence at wider spatial scales on focal incidence indicate high epidemiological connectivity. These effects are driven in part by vaccination at these spatial scales, either through reduced incidence in other vaccinated villages lowering the risk of rabid dogs roaming into the focal village or from dog owners in the focal village accessing vaccination campaigns in neighbouring villages. Therefore, high coverage and resulting low incidence elsewhere mitigate local coverage gaps. In contrast, high coverage in a small area has only limited impact if the rest of the population is poorly vaccinated, consistent with the findings of previous simulation studies[38,63]. This interplay between dog vaccination and movement may have complex impacts on incidence that are also influenced by population configuration. Studies from Serengeti district [64] and Bali, Indonesia [65] identified impacts of prior incidence in nearby areas on rabies occurrence. Unlike those studies, we did not detect decreased impacts with distance from the focal village, possibly because we studied only the collective impacts of non-bordering versus bordering villages (80+ vs ∼5); evaluating individual village impacts would be more likely to reveal distance-related effects. The observation that logging prior incidence (such that the rate of increase in focal incidence diminishes with increasing prior incidence) improves model fit is possibly a consequence of susceptible depletion, or local responses to outbreaks.

We anticipated that including prior incidence in our model would mask the effects of vaccination coverage at different spatial scales. This was confirmed in our results and was expected because prior incidence is partly driven by prior coverage. The strong effect of prior incidence, and the reduction in the quality of fit when prior incidence is removed from models are also unsurprising. Prior incidence, in addition to incorporating part of the effect of prior vaccination, also incorporates information about actual transmission events that drive the force of infection coming both from within the district and from incursions. Difficulties in separating out the impacts of correlated variables is a limitation of the GLMM framework. A mechanistic transmission model could more explicitly explore impacts of spatially heterogeneous vaccination, as per simulation studies [36,38,48]. However, this approach requires estimation of (or assumptions about) many parameters, and is vulnerable to assumptions about the important mechanisms to be included. For these reasons, we explored GLMMs with and without the effect of prior incidence, to produce models both with a high quality of fit and from which we could extract independent effects of vaccination.

We inferred an average of 7 incursions annually, with incursions more frequent near district borders. Incursions in the district centre likely result from human-mediated transport, which is an increasingly recognised problem [33,36,38,66,67]. Our inference methods have limitations; local transmission could be misidentified as incursions due to unobserved transmission, unusually long incubation periods or dog movements, while real incursions could go undetected within existing foci. We expect less misidentification at low incidence, suggesting inferred incursions should have increased post-2018. However, no increase was observed, possibly because real incursions decreased from the indirect effects of vaccination on circulation in neighbouring districts and/or the direct impact of vaccination in those districts from 2020 [30,68]. Misidentification may contribute to fewer inferred incursions in the district centre, where incidence was generally higher. Simulation studies could validate the performance of incursion assignment, and incorporating viral genomes may improve accuracy [33]. Regardless, the proportion of cases identified as incursions is far higher in later years (>20% of cases since 2019, reaching as high as 50% in 2022). This encouraging indication that local transmission has been interrupted, also serves as a warning that incursions are now a major driver of continued transmission and, no matter how well district-level control measures are implemented, elimination will require expanded vaccination. The example of Latin America shows how scaled up vaccination has largely eliminated dog-mediated rabies, with a contracting set of foci remaining in only the most challenging settings [22,40]. In Tanzania, rabies vaccination in Serengeti district began to extend across the surrounding Mara region in late 2020 [30,68]. Estimated incursions did not decline in 2021-2022, but this may change as rabies is controlled in neighbouring districts.

A limitation of our rolling coverage estimates is the assumption that all dogs are equally likely to be vaccinated in each annual campaign. In reality, dog owners who chose not to attend a previous campaign may have had reasons (e.g. difficulty in handling dogs, distance to the vaccination point [57]) that make them likely to continue not to bring their dogs in subsequent years. Alternatively, attendees from previous campaigns may be less inclined to return, assuming their dogs are already protected. Given little data on these two behaviours, and evidence that repeat campaigns attract a mix of new and previously vaccinated dogs [25], we assumed the behaviours cancel out. However, this may have led to bias in our estimates. When calculating both campaign and rolling coverage, we also assume that the same dog is not vaccinated more than once a year. Campaigns were typically held for one day each year in each village, with rare follow-up vaccinations in response to outbreaks targeting unvaccinated dogs, and we expect owners are unlikely to bring their dogs to multiple campaigns in different villages in a year. However, owners could unknowingly revaccinate dogs that they recently acquired from another village, leading to overestimation of coverage.

A number of our results warrant further investigation. We consistently found a positive effect of human:dog ratio and negative effect of dog density on rabies incidence. We hypothesise that settings with higher human:dog ratios have less awareness of dog behaviour, leading to rabies circulating with less intervention, while higher dog densities mean more cases are observed at the same incidence, leading to greater visibility and faster intervention. Studies of dog ownership practices in different settings and of how human responses to rabid dogs change based on recent incidence may increase understanding of these effects. We also observed that human rabies exposures from jackals and other wildlife were more common than by livestock, despite livestock comprising a higher proportion of animal cases than wildlife. There are multiple potential explanations for this; rabid livestock may be less aggressive and their bites easier to avoid, or wildlife cases that do not cause exposures less likely to be observed and thus under-represented in case data (compared to economically valued livestock).

Returning to our three initial hypotheses for why rabies continues to circulate despite control efforts, we conclude that: 1) spatial heterogeneity in dog vaccination was detrimental for control, and vaccination was only effective in interrupting endemic circulation when implemented consistently, underlining the importance of monitoring fine-scale coverage to promptly identify and address gaps; 2) mean coverage over the district was relatively low, but once heterogeneity was addressed, rabies was controlled nonetheless – a promising result given hard-to-reach populations with fast demographic rates; 3) Incursions from unvaccinated areas outside the district are frequent and now represent a major source of residual transmission. High epidemiological connectivity within the district, and frequent incursions from outside, interact with heterogeneous vaccination in complex ways that should be considered in programme design. It is now vital that dog vaccination efforts be scaled up to ensure that impacts are sustained, to maximise their benefit to all, and to achieve the ‘Zero by 30’ goal.

## Materials and Methods

### Dog vaccination

We focus on the epidemiological dynamics of rabies in Serengeti District, northwest Tanzania from January 2002 to December 2022 using contact tracing and demographic data over the same period, and mass dog vaccination data with associated population estimates from January 2000 to December 2022 (to establish the level of pre-existing vaccination before the main study period). Between October 1996 and February 2001, four vaccination campaigns were carried out in Serengeti District [25]. These campaigns covered all the villages in the district as it was then, but we note that the district has since increased in area, with new villages incorporated that were previously part of the adjacent district, Musoma, and that were not covered in those earlier campaigns. Following the last of these four campaigns, a household survey estimated 73.7% coverage in the district. There was no vaccination in Serengeti district in 2002, but in 2003 Rabies Free Africa initiated annual campaigns in the villages in the East and South of the district, aiming to protect people and their animals and to prevent rabies spilling over into vulnerable carnivore populations in Serengeti National Park [42]. In 2004, vaccination campaigns were expanded to more villages with support from the District Veterinary Office.

Since then dog vaccination typically takes place annually using a central point strategy where dogs are brought to a central location, such as a village centre, church, or school in each village. A vehicle equipped with a loudspeaker is used to advertise vaccination the day before the campaign. Posters advertising the vaccination day are hung at busy locations within each village, such as village headquarters office, dispensaries, schools, churches and mosques. On vaccination days, dogs are registered, recording owner name, dog name, age, and sex. Rabies vaccination is offered free of charge using the Nobivac Rabies vaccine (MSD Animal Health, Boxmeer, The Netherlands). Dog owners receive a vaccination certificate for each vaccinated dog.

Since 2003, dogs have occasionally been vaccinated outside of central point campaigns in response to localised transmission. Since late 2020, villages in the northwest of the district have been part of a large-scale vaccination trial across Mara Region. A subset of those villages are following a continuous strategy, where vaccinations happen year-round [30] while the remaining villages continue annual central-point campaigns. Date, village and number of dogs vaccinated are available for all described vaccination events throughout the study and can be found in our Github repository: https://github.com/boydorr/Serengeti_vaccination_impacts.

### Rabies incidence and exposures

Contact tracing was carried out following the methods outlined in Hampson et al. [44] This approach generated data on 3,973 probable animal rabies cases – of which 3,362 were domestic dogs, 368 were livestock, 159 were wildlife, 81 were domestic cats, and three did not include a record of species – and 1,612 probable human rabies exposures in the years 2002-2022. It has previously been estimated that these methods identify 83-95% of carnivore cases in the district [69]. Each probable animal case and probable human exposure was georeferenced and time-stamped, with the identity of the biting animal recorded where possible.

### Dog population estimation

Human population counts were available at the village level from the 2012 government census [70], and at ward-level for the 2002 and 2022 censuses [71]. Village-level estimates for 2002 and 2022 were obtained by assuming that the population in each ward was divided between the villages in that ward in the same proportions as in 2012. For each village *v*, we then estimated the human population *H* for every month *m* in the period from January 2000 (i.e. *m*=1) to December 2022 (i.e. *m*=276) to be:

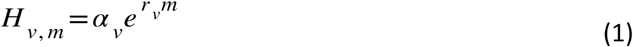

where parameters *ɑ_v_* and *r_v_* were estimated for each village using the three census populations by non-linear least squares.

A georeferenced household census of humans and dogs in Serengeti District was completed between 2008 and 2016 [41], with each village censused once at some point during this period. This allowed estimation of village-specific human:dog ratios *R_v_*. The dog population in every village each month was estimated as:

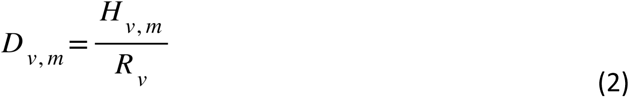

We also mapped the dog population to a grid of 1km^2^ cells by first assigning each cell to a village within Serengeti district. For each village, the dog population in each month (*D_v,m_*) was then distributed among cells of that village in proportion to the number of dogs in those cells at the time of the georeferenced census.

### Vaccination coverage

We calculated the numbers of dogs vaccinated in each village in each month in 2000-2022, *V_v,m_*. To allow estimation of existing levels of vaccination at the time contact tracing began in 2002, we included information about the final vaccination campaign in the district prior to 2002, which was completed in 2000-2001 [25]. Monthly village-level data were not available for this campaign, so we assumed that the 73.7% of dogs estimated to have been vaccinated by post-vaccination household surveys (completed within two days of the vaccination campaign in each village) were distributed evenly across the campaign months (May 2000 to February 2001) in all villages that were part of the district at that time.

Village campaign coverages each year *y* (Fig. 2E) were calculated as:

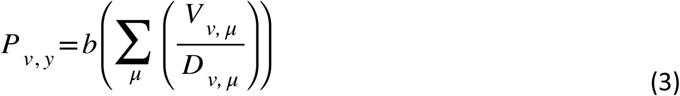

where *μ* are the months of *y*. This assumes that no dog is vaccinated twice within the same year. To prevent estimates of *P_v,y_*>1 (e.g. due to dog populations being underestimated or dogs being taken for vaccination outside their home village), we use a generalised sigmoidal bounding function *b*(*x*):

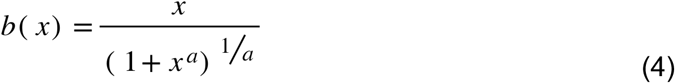

where we set *a*=6. This means that values of *x*<0.7 experience little change, while *x* =1.0 would be reduced to a more realistic 0.89 (Fig. S14).

Annual district campaign coverages were calculated as:

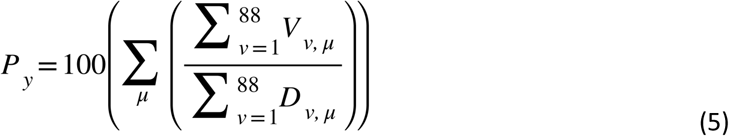

When estimating monthly village-level rolling vaccination coverages (i.e. the proportion of dogs in the population that have been vaccinated at least once since January 2000), we assume that numbers of vaccinated dogs decline geometrically each month with ratio:

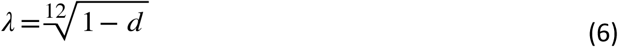

where *d*=0.448 is the proportion of dogs that die in a year [72]. Assuming that all dogs are equally likely to be vaccinated each year, regardless of previous vaccination status, and that the same dog cannot be vaccinated twice in the same year, the number of vaccinated dogs in a village in a given month can be estimated as

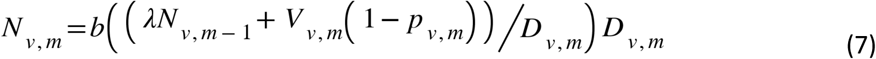

where *p_v,m_* is the proportion of dogs that are available to be vaccinated this month (i.e. not already vaccinated in the current year) that had already been vaccinated in a previous year:

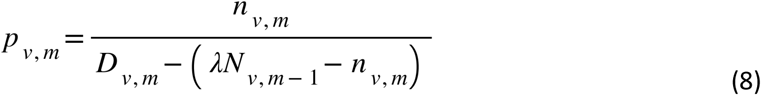

and *n_v,m_* is the number of vaccinated dogs that were not vaccinated in the current year:

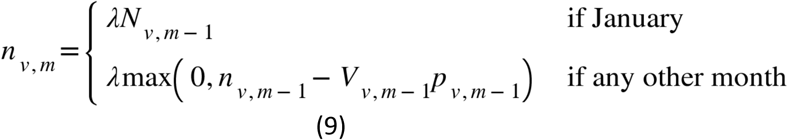

Finally, we obtained the rolling vaccination coverage for every village and month by:

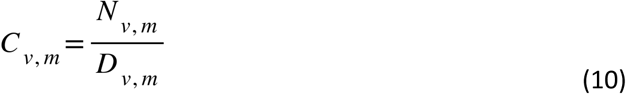

and rolling vaccination coverage over the district in each month by:

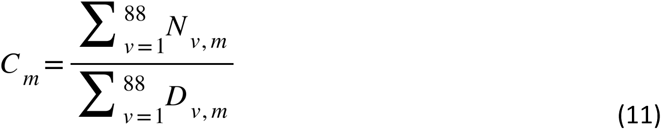

The level of heterogeneity in rolling coverage over the district each month *H_m_* was quantified using the population-weighted standard deviation in village rolling coverage:

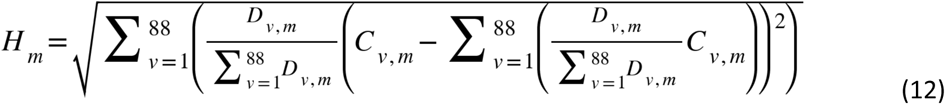

### Modelling the impact of vaccination on rabies incidence

To investigate the impact of vaccination and other variables on rabies incidence, we developed a univariate negative binomial generalised linear mixed model (GLMM), where the single response variable was the number of probable dog rabies cases in a village in a given month. An offset equal to the log of the estimated dog population *D_v,m_* was included, allowing us to model incidence as cases per dog. Village was included as a random effect. Multiple fixed effects (each described in full below) were included together in the model including: 1) either rolling vaccination coverage or susceptibility (1 – rolling coverage) in the focal village over the prior two months; 2) either arithmetic mean coverage or power mean susceptibility over villages bordering the focal village over the prior two months; 3) either arithmetic mean coverage or power mean susceptibility over villages not bordering the focal village over the prior two months; 4) incidence in the focal village over the prior two months; 5) incidence over villages bordering the focal village over the prior two months; 6) incidence over villages not bordering the focal village over the prior two months; 7) dog density in the focal village in the current month; 8) the human:dog ratio in the focal village.

All analyses were conducted in R [73], with models fitted using Stan [74], via the RStan package [75] for models that included power means of susceptibility to allow estimation of both the power parameter *p* (equation (13)) and the model coefficients and otherwise via the brms package [76]. For all variable coefficients we assumed a normal prior, N(μ=0, σ=100,000). For the size parameter of the negative binomial distribution, we use a gamma prior, G(ɑ=0.01, β=0.01) (the default prior selected for this parameter in brms). An exponential prior, Exp(λ=0.001), was used for the standard deviation of the village random effect. To discourage what we believed to be unrealistically large effects of heterogeneous coverage (Fig. S6), we used a normal prior for *p* centred on the arithmetic mean with a standard deviation of two, N(μ=1, σ=2). For models fitted in brms, we checked assumptions using simulated residuals via the DHARMa package [77].

Since >90% of rabies incubation periods recorded from contact tracing were less than two months, we include mean estimated vaccination coverage in the village *v* over the prior two months (*C_v,m-1_+C_v,m-2_*)/2 as a fixed effect. As we also wanted to explore impacts of conditions in areas outside the focal village on focal incidence, we included fixed effects of mean vaccination coverage at two wider scales over the prior two months. The first of these was at the borders of the focal village. We obtained this variable by identifying all villages that shared a border with the focal village and calculating the proportion of the border shared with each of these villages. Bordering vaccination coverage in a given month was then calculated by multiplying these border proportions by vaccination coverages in these bordering villages, and summing over the bordering villages, i.e. the border-weighted arithmetic mean coverage. Vaccination coverage at borders with Serengeti National Park was assumed to be equal to the average of the vaccination coverages in the other bordering villages, and coverage at borders with the rest of Mara District was set to 9% (the coverage estimated in Serengeti District prior to mass vaccination campaigns [25]). Finally, we introduced a fixed effect of the prior population-weighted arithmetic mean vaccination coverage over the dog populations of the rest of the district villages not sharing a border with the focal village.

To explore the impact of heterogeneity in coverage within the district on incidence in focal villages, we also considered a version of this model where the arithmetic mean coverages over bordering and non-bordering villages were replaced by power mean susceptibility (where susceptibility=1-coverage) at these scales. A power mean *M_p_* over a sample, is calculated as:

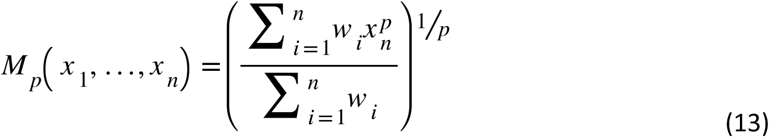

where *w_i_* are the weights applied to each observation (length of shared border for mean susceptibility of bordering villages, and dog population for mean susceptibility of non-bordering villages). When fitting this model, we estimated the value of *p* in addition to the two GLMM coefficients for power mean susceptibility in bordering and non-bordering villages. We focus on power means of susceptibility rather than of vaccination coverage to enable calculation of logarithms of the power mean elements (to allow first-order approximation when *p* is close to zero and to provide stabilisation at large *p*); vaccination coverages of zero are plausible and occur frequently in the data, while susceptibilities of zero are both implausible in this setting, and corrected by equation (4).

Current incidence of dog rabies was expected to be highly dependent on prior incidence, so we included an effect of mean incidence in the focal village over the prior two months. Prior incidence at the wider spatial scales was included as described for vaccination. Monthly incidence at borders with the rest of Mara Region was assumed to be 0.01/12, where 12 represents the number of months in the year and ∼0.01 was the highest proportion of Serengeti’s dog population infected in any given year1. This assumption is based on the rest of Mara Region being largely unvaccinated, and thus having generally high incidence. Incidence at the borders with Serengeti National Park was assumed to be equal to the average incidence in the other villages bordering a focal village. We compared a model where the prior incidence variables at each of the three spatial scales were logged with one where they were unlogged, selecting the best based on WAIC. We also explored models where some of all of the prior incidence variables (which correlate with prior coverage/susceptibility) were removed (as outlined in Table 1), so as to estimate independent effects of coverage/susceptibility.

We incorporated fixed effects of the log of dog density in the village and the human:dog ratio *R_v_*. When calculating dog density in a village we used the number of 1km^2^ grid cells assigned to the village that were occupied by at least one household during the Serengeti dog census as the denominator. This denominator was chosen to account for the fact that many villages, particularly on the North and South borders (Fig. 1A), have large unoccupied areas that would otherwise bias density estimates.

In addition to the models described above, which use data at relatively fine spatiotemporal scales (village and month), we investigated the impact of aggregating data by fitting models at the remaining three combinations of village/district and monthly/annual scales. Prior distributions remained as previously specified. At the monthly, district scale, a univariate negative binomial generalised linear model (GLM) was fitted, where the single response variable was cases in the district each month, with an offset of the logged district dog population. Multiple fixed effect combinations were considered as outlined in Table S2. These fixed effects included either rolling vaccination coverage at the district level (equation (11)) or power mean susceptibility calculated over all 88 villages using equation (13), both averaged over the prior two months. Monthly rabies incidence in the district was calculated by dividing the total cases that month by the estimated total district dog population that month, and again incorporated in the model as an average over the prior two months. District-level dog density was included by dividing the total district dog population each month by the number of occupied 1km^2^ grid cells in the district.

The third set of models at the annual, village scale, was fitted as univariate GLMMs with a response of the number of cases in a village in each year, with an offset of the logged mean village dog population that year. Prior vaccination at the three spatial scales (village, borders, rest of district) was incorporated in the form of the campaign coverage *P_v,y_* (equation (3)). We fitted models with fixed effects of: 1. *P_v,y_* last year; 2. Mean *P_v,y_* over the last two years; 3. Mean *P_v,y_* over the last three years; 4. *P_v,y_* last year and logged incidence last year. Log dog density and human:dog ratio in the focal village were also included in all four models, with village as a random effect. All fixed effect combinations are described in Table S3.

Finally, we fitted a fourth set of models at the annual, district scale that were univariate GLMs with a response of cases in the district each year and an offset of the logged mean district dog population over that year. The campaign coverage in the district each year *P_y_* (equation (5)) was included in the same four combinations as for *P_v,y_* in the annual village-level models. All of these models included log dog density as a fixed effect (see Table S4 for details).

Annual models included fixed effects of campaign coverage rather than rolling coverage to make results comparable to a previous study of impacts of vaccination on rabies incidence [39]. The importance of dogs being carried over from previous campaigns was instead considered by our exploration of models that averaged prior campaign coverage over the previous 2-3 years.

### Estimating incursions

To identify cases that likely originated from transmission outside Serengeti District (i.e. incursions), we reconstructed transmission trees using the treerabid package in R [69,78]. For this analysis, we excluded cases in herbivores, since these are unlikely to cause onward transmission, reducing the analysed cases to 3,600. Tree reconstruction required distributions for the serial interval *S* (the time between onset of symptoms in an offspring case and in its parent case) and dispersal kernel *K* (the distance between cases and contacts regardless of whether these developed rabies). We calculated serial intervals from 1,156 paired dates of symptoms onset from cases and their recorded rabid biter and Euclidean distances between locations of 6,897 contacts and their recorded biter. For both *S* and *K* we fitted gamma, lognormal and Weibull distributions using the fitdistrplus package [79], with the best-fitting distributions - lognormal for *S* and Weibull for *K* - chosen based on AIC (Table S5).

While fitting these distributions, censoring was applied for distances as described previously [69]. Distances of <100m where the biter was a dog with known owner, and an accurate home location, were interval censored between 0 and 100m to account for homestead sizes. For unknown dogs and wildlife, where locations were recorded for first observation rather than home location, the true distance to contacts was assumed unknown (right censored), but with a minimum of the recorded value or 100m, whichever was larger, on the basis that biters from <100m away would be recognised. Parameters of the fitted distributions (Table S5) have changed little from those estimated previously [69], despite an additional seven years of contact tracing. A comparison of the data and best-fitting distributions is shown in Fig. S13.

The transmission tree reconstruction algorithm [78] was used to probabilistically assign a progenitor to each case *i* with no known rabid biter (65.75% of all carnivore cases). To be considered a possible progenitor of *i*, a case *j* has to have a symptoms onset date preceding that of *i*, and the serial interval and distance between *i* and *j* must be smaller than a selected quantile of the distributions *S* and *K*. This quantile, known as the pruning threshold, is chosen to prevent assignment of progenitors that are unlikely to be correct given the separation of cases in space or time. We explored pruning thresholds of 0.95, 0.975, and 0.99; the maximum serial intervals and distances defined by these thresholds are given in Table S5. The probability of each case j∈{1,…,n} that meets these pruning criteria being randomly selected as the progenitor of *i* is:

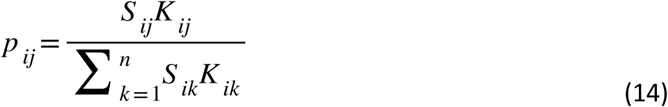

where *Sij* and *K_ij_* are the probabilities of the serial interval and distance between *i* and *j* based on reference distributions *S* and *K*. In cases where *n*=0, no biter was assigned, with *i* being assumed to be an incursion. To account for recorded uncertainties in the timing of bites and symptoms onset, 1,000 bootstrapped datasets were generated by drawing dates randomly from a uniform distribution over the recorded uncertainty window, while preserving the correct sequence of events between known parent and offspring cases. We used the best-fitting lognormal distribution for *S* (Table S5) and when *i* was a dog with known owner and accurate location we used the best-fitting Weibull distribution for *K*. When *i* was an unknown dog or wildlife, where the recorded location was generally the location *i* was observed biting, not *i*‘s home location, *K* was a Weibull distribution fitted to simulations of the Euclidean distance travelled as a result of two draws from the previously fitted Weibull with a uniform random change in direction between them (representing the movement of *j* to *i*, followed by the movement of *i* to its recorded location). For each pruning threshold, we identified probable incursions as those cases identified as an incursion more frequently than they were assigned to any other progenitor within the set of 1,000 bootstrapped trees. Inferred incursions and proportion incursions were notably high in 2002, as an expected artefact of beginning contact tracing, and are thus not considered in the results. Progenitors of some 2002 cases likely occurred in 2001, and thus would have gone undetected. Additionally, detection was likely lower in 2002 as methods were still being honed.

### Data and code availability

All analyses were carried out using the R statistical computing language, version 4.2.0 [73]. Deidentified data and code files are located in our Github repository: https://github.com/boydorr/Serengeti_vaccination_impacts.

### Ethical permissions

Ethical approval for this research was obtained from the Tanzania Commission for Science and Technology, the Institutional Review Boards of the National Institute for Medical Research in Tanzania and of Ifakara Health Institute, and the Ministry of Regional Administration and Local Government (NIMR/HQ/R.8a/vol.IX/300, NIMR/HQ/R.8a/vol.IX/994, NIMR/HQ/R.8a/vol.IX/2109, NIMR/HQ/ R.8a/vol.IX/2788, and IHI/IRB/No:22-2014).

## Acknowledgements

We are grateful to the local communities in Serengeti District and to government staff from the animal and public health sectors for their ongoing support. We also thank Rabies Free Africa for their dog vaccination efforts, MSD Animal Health for donating vaccines for the vaccination campaigns, Machunde Bigambo and Matthias Magoto for supporting data collection, and Daniel Haydon for providing constructive feedback that greatly improved this work.

## Funding

This work was funded by the Wellcome Trust (grants 095787/Z/11/Z, 207569/Z/17/Z and 224520/Z/21/Z to KH, https://wellcome.org/) and by the Department of Health and Human Services of the National Institutes of Health (grant R01AI141712 to FL and KH, https://www.nih.gov/). The funders had no role in the study design, data collection and analysis, decision to publish, or preparation of the manuscript. The content is solely the responsibility of the authors and does not necessarily represent the official views of the National Institutes of Health.

### Author Contributions

**Elaine A Ferguson:** Conceptualization, Data curation, Formal analysis, Methodology, Software, Validation, Visualization, Writing – original draft, Writing – review and editing. **Ahmed Lugelo**: Investigation. **Anna Czupryna**: Investigation, Writing – original draft, Writing – review and editing. **Danni Anderson:** Data curation. **Felix Lankester:** Funding acquisition, Project administration, Writing – review and editing. **Lwitiko Sikana:** Investigation. **Jonathan Dushoff:** Formal analysis, Methodology, Supervision, Writing – review and editing. **Katie Hampson:** Conceptualization, Funding acquisition, Methodology, Project administration, Supervision, Writing – original draft, Writing – review and editing.

## Supplementary information

### Supplementary Videos

**<u>Video S1</u>**

**Video S1: Monthly spatial distribution of dog cases, human exposures and dog vaccination coverage over the study period.** Estimated vaccination coverage at the village-level for each month in 2002-2022 is indicated by the colour scale. Locations of dog cases (blue points), human exposures (purple triangles) and human deaths (large black triangles) each month are indicated.

### Supplementary Figures

**Figure S1.**
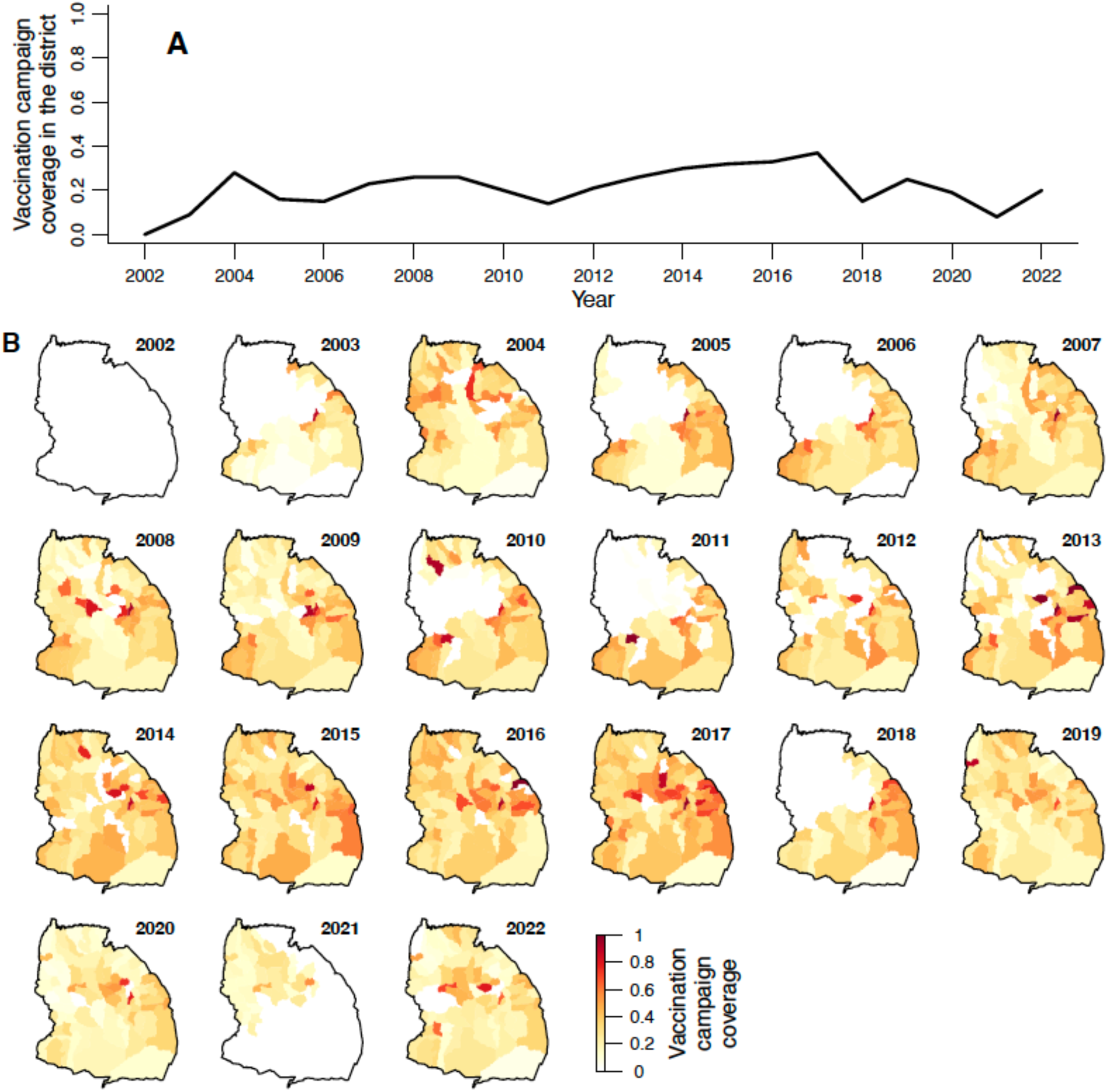
Campaign vaccination coverage each year from. 2002-2022 A) at district level and B) at village level.

**Figure S2.**
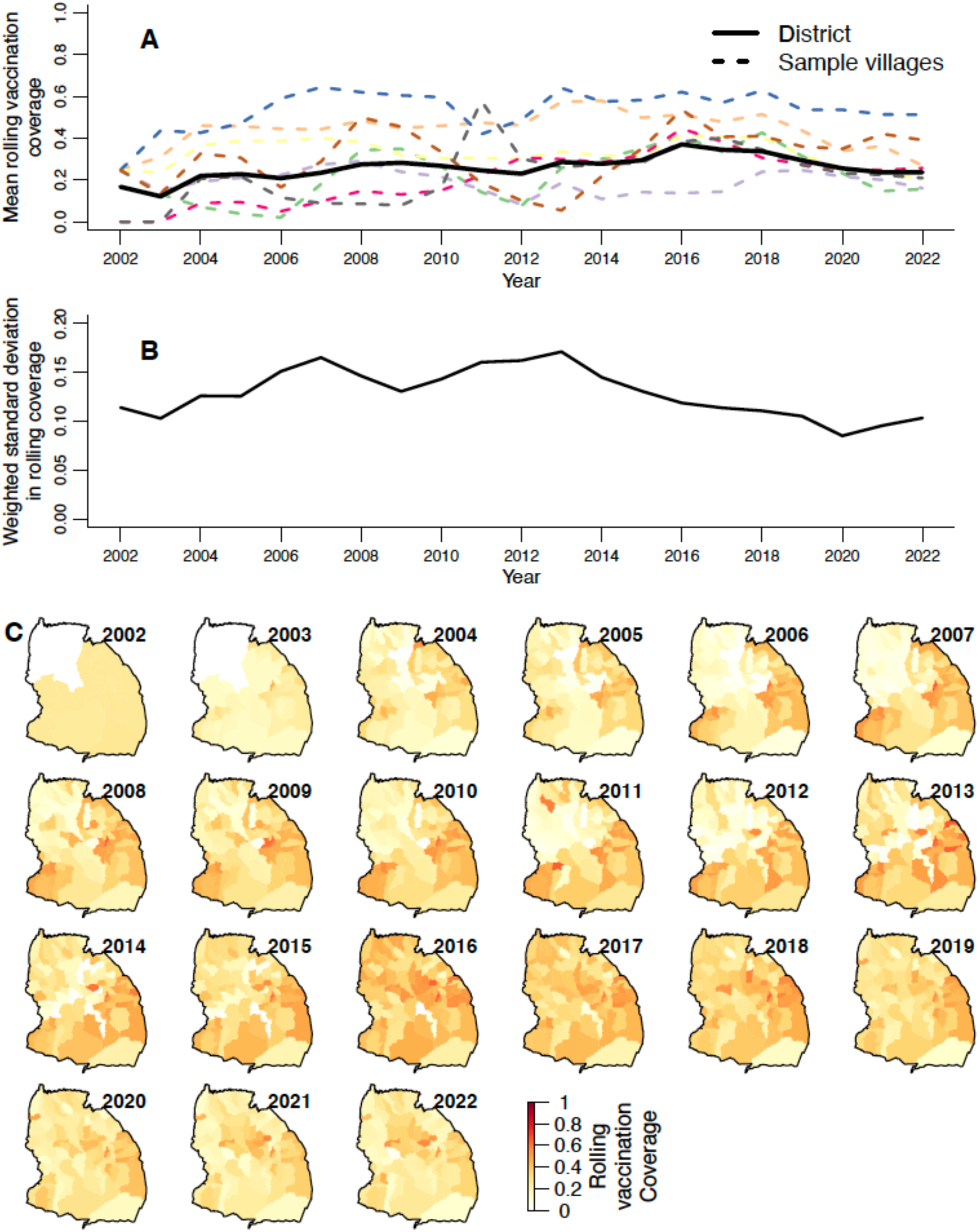
Mean of and spatial heterogeneity in rolling vaccination coverage each year. A) The mean rolling vaccination coverage in Serengeti District over each year (12-month averages of the values in Fig. 2C) is indicated by the solid black line. Mean rolling coverages each year for 8 randomly selected villages are indicated by dashed coloured lines. B) The weighted standard deviation in the rolling vaccination coverage over the villages in Serengeti District (yearly means of the values in Fig. 2C) . C) Mean rolling vaccination coverage in each village over each year from 2002-2022 is indicated by the colour scale.

**Figure S3:**
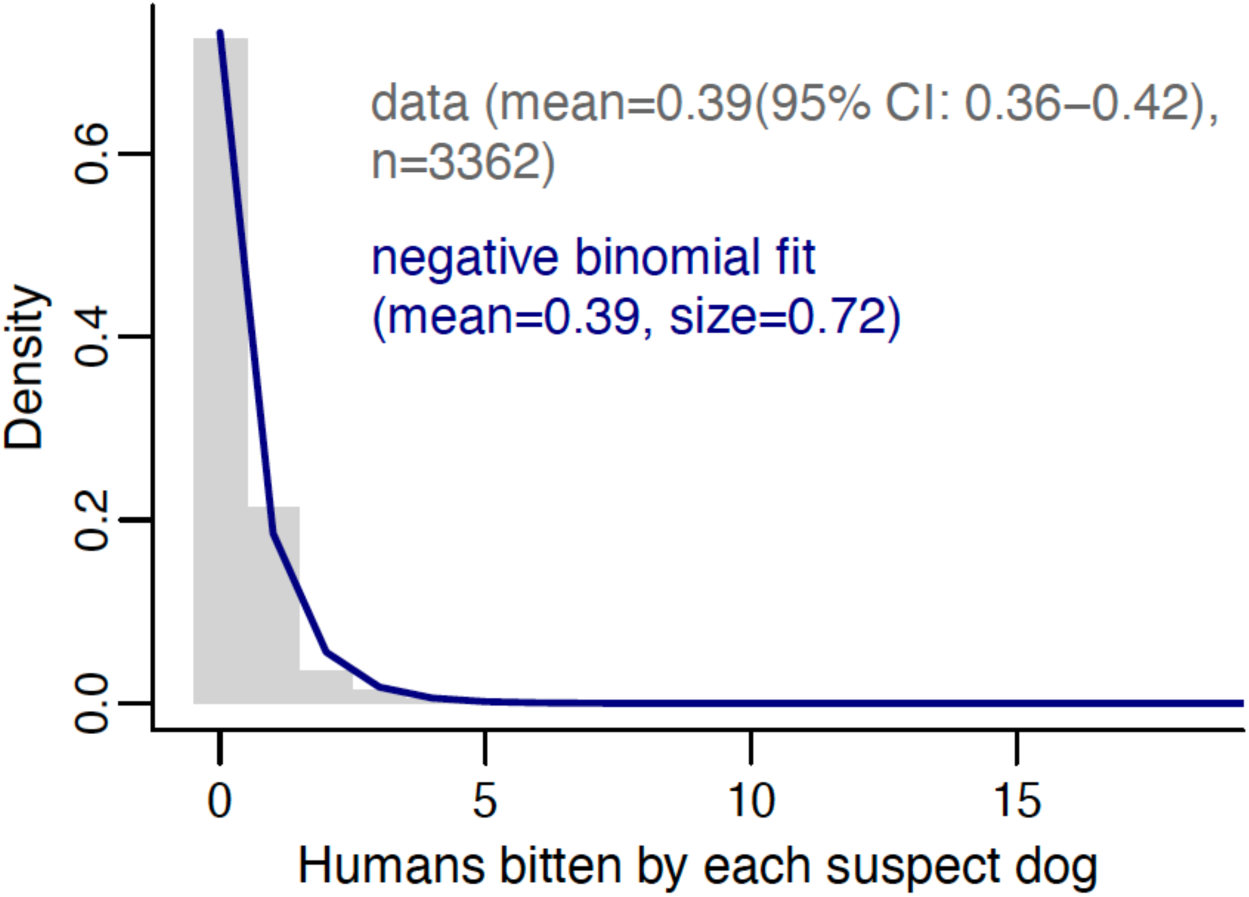
Human exposures per rabid dog. Histogram of human rabies exposures by each rabid dog from contact tracing data (grey bars), with fitted negative binomial distribution (blue line).

**Figure S4:**
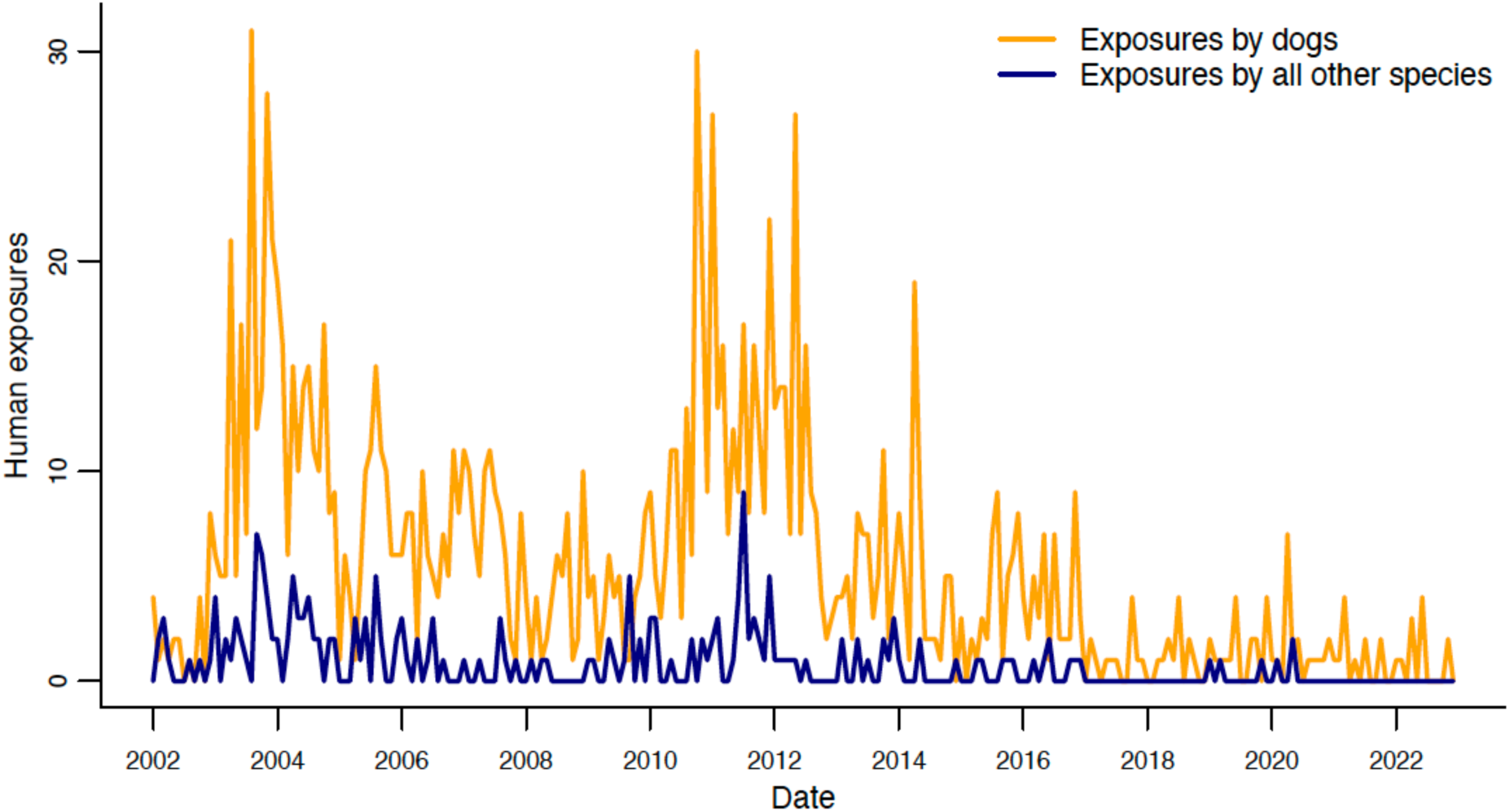
Probable human exposures by dogs vs. by all other species from contact tracing.

**Figure S5:**
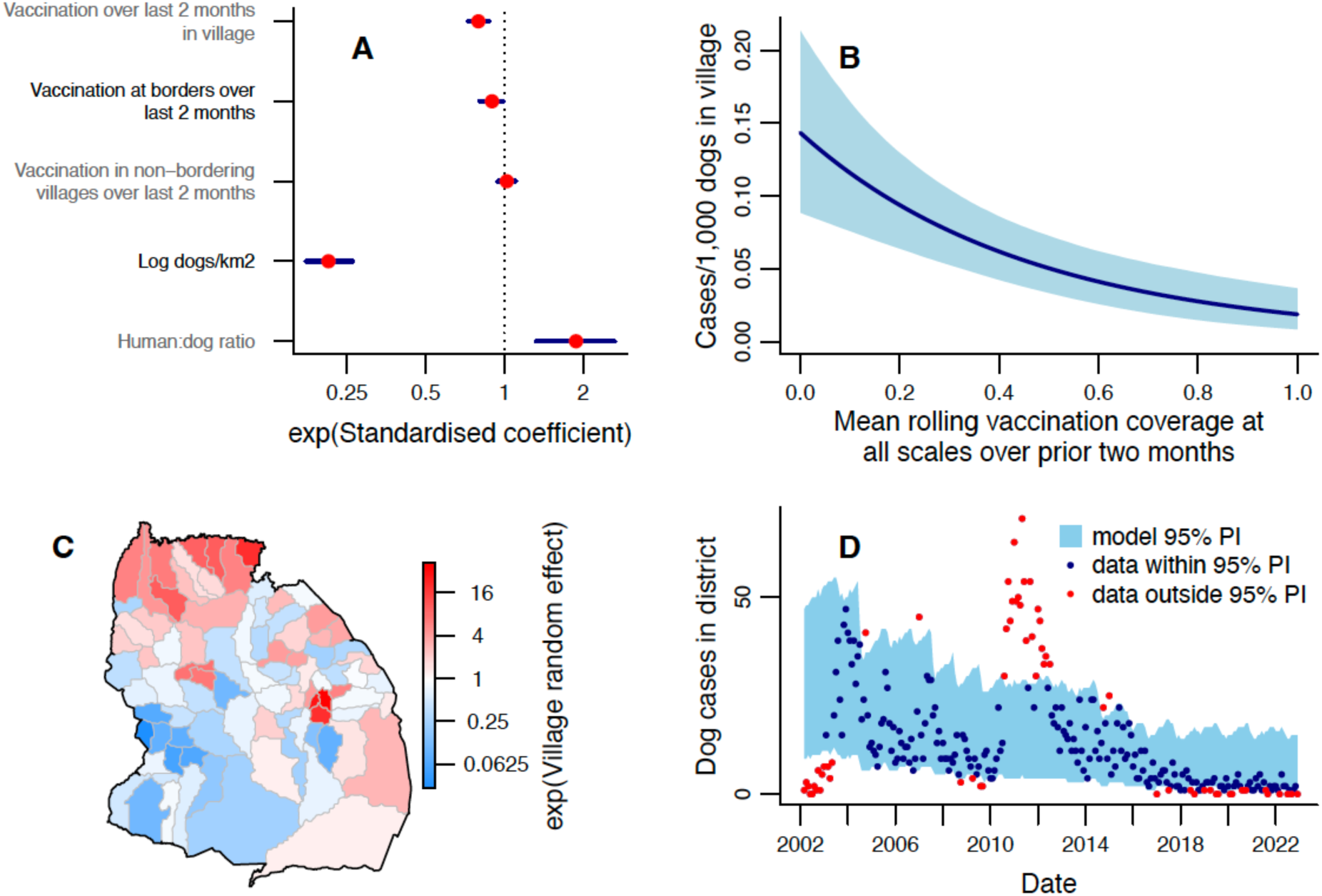
Impact of removing prior cases/dog explanatory variables from the monthly village-level GLMM for current cases per dog. (compare with Fig. 3). A) Exponentiated standardised values of the coefficients estimated for each explanatory variable, with 95% CrIs. B) Line shows the expected cases/1,000 dogs (number of dog cases normalised by dog population) in a village this month for different mean rolling vaccination coverages across the focal village and district in the prior 2 months. Shaded areas show 95% credible intervals (CrIs), and predictions were obtained using average values of unspecified explanatory variables. C) Exponentiated random effect values for each village in the district. D) Comparison of observed monthly dog cases (points) with the 95% prediction interval from the fitted model. Data points in red fall outside the 95% prediction interval (PI).

**Figure S6:**
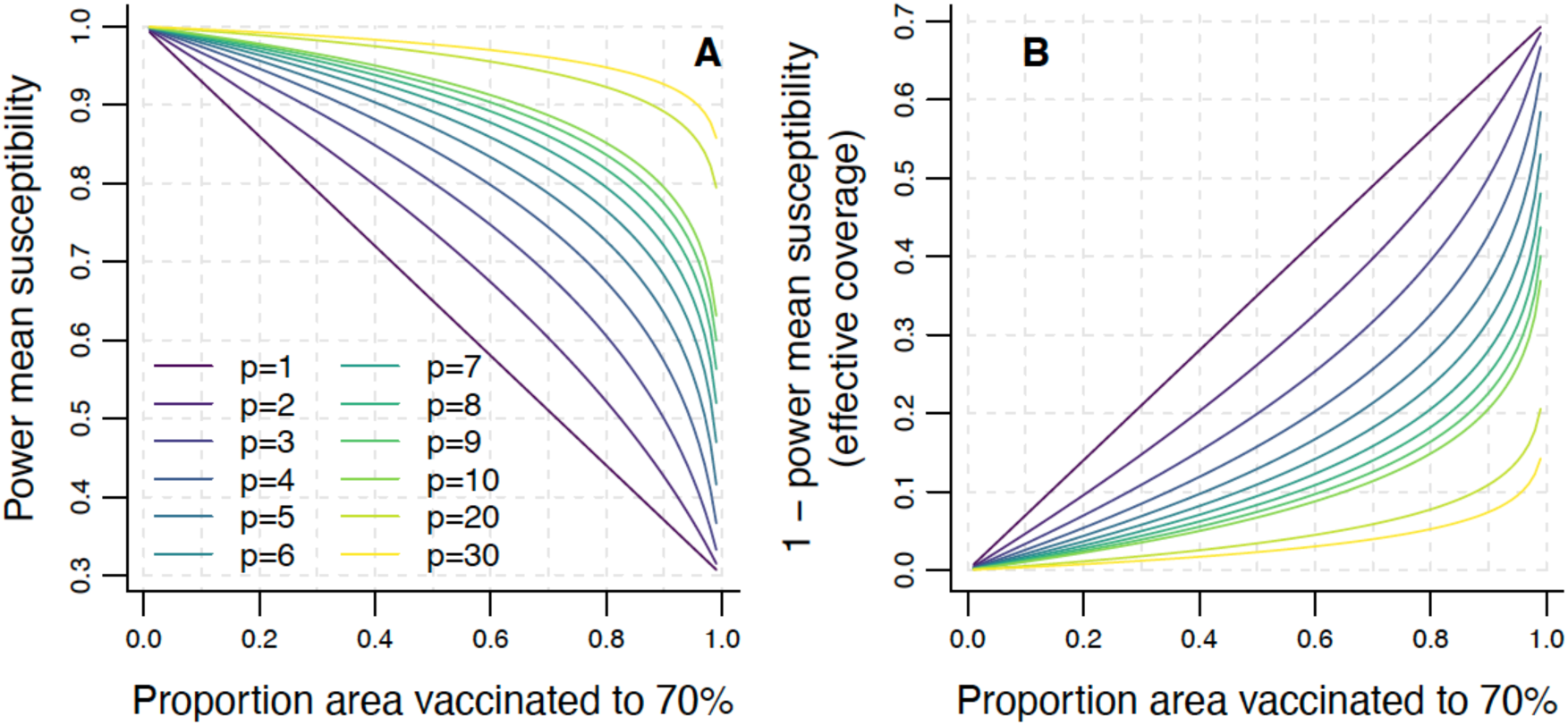
The impact of different powers on the power mean of susceptibility under different levels of heterogeneous vaccination. Here we assume a landscape where an increasing proportion of the area is vaccinated to 70% (30% susceptibility) while the remaining proportion remains at 0% coverage (100% susceptibility). We then calculate A) the power mean susceptibility and B) the effective coverage (1 – power mean susceptibility) at each proportion of area vaccinated at a range of values of the power *p* (equation (13)). *p*=1 is the arithmetic mean, and represents a scenario where, for a given proportion of dogs being vaccinated, the effective level of vaccination is the same regardless of how these vaccinated dogs are distributed over the area, i.e. heterogeneity in vaccination does not reduce (or increase) the impact of that vaccination on rabies cases. We used these curves showing the impact of different powers to select the prior distribution for *p*∼N(μ=1, σ=2). If 99% of the area is covered, then the arithmetic mean coverage is 69.3%. If *p*=2, then the effective coverage for the heterogeneous landscape is 68.5%; 0.8% lower than if vaccination had been homogeneous. If *p*=5, however, effective coverage would be 58.4%, which is 10.9% below the arithmetic mean, despite only 1% of the area being uncovered, which seems an excessively large effect. The choice of σ=2 was therefore made to exclude *p*≥5 from the a priori confidence interval.

**Figure S7:**
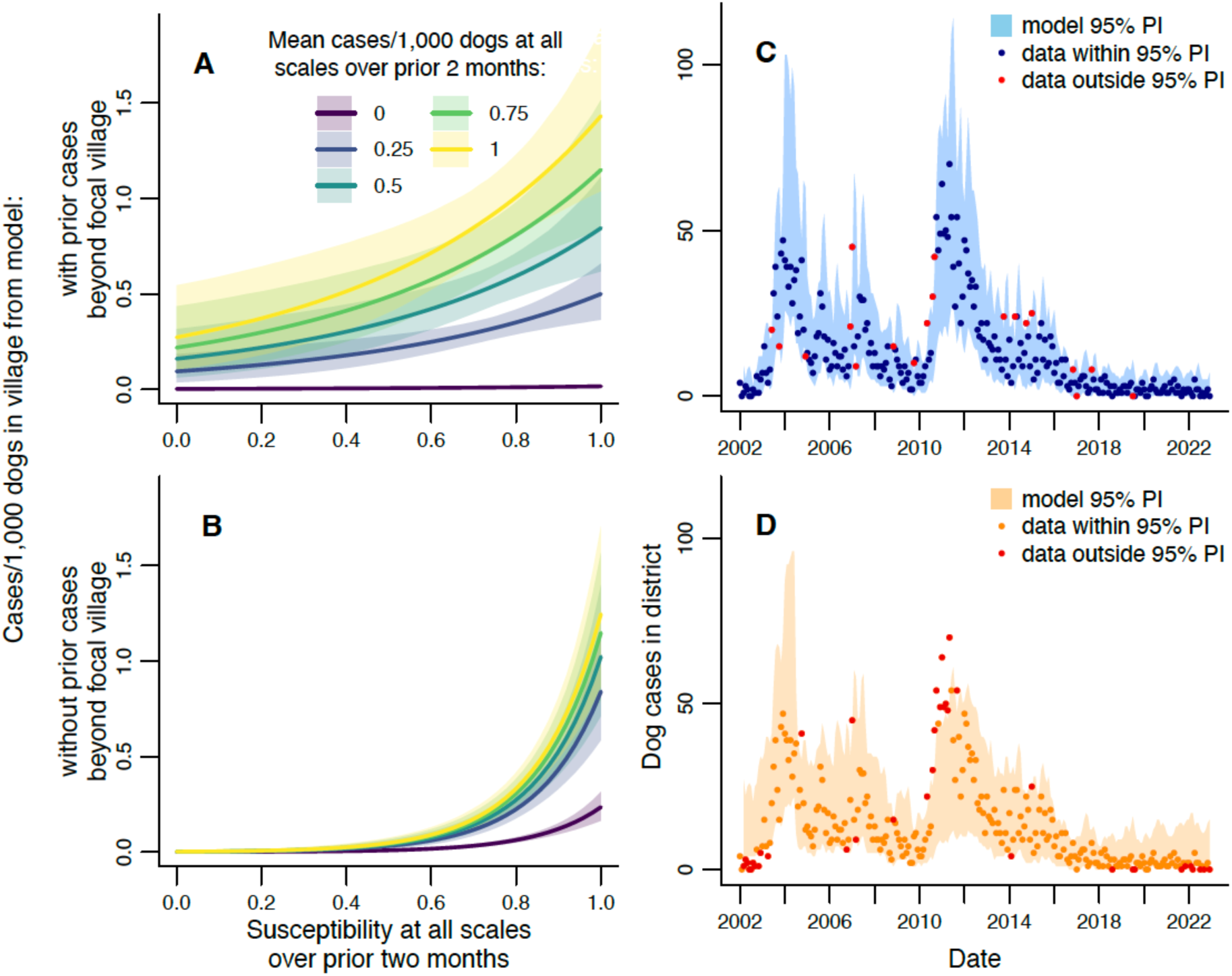
Impact of power mean susceptibility on rabies incidence at the village level and quality of model fits to data. A-B) Expected cases/1,000 dogs in a village from models with (A) and without (B) effects of prior incidence beyond the village. Predictions are shown for different mean susceptibilities (assuming homogeneous vaccination, i.e. power mean susceptibilities beyond the village equal susceptibility in the village) and mean cases/dog in the prior 2 months. Prior cases/dog values represent the observed district-level range and shaded areas show 95% CrIs, Predictions were obtained using average values of unspecified explanatory variables. C-D) Comparison of observed monthly dog cases (points) with the 95% prediction interval from the fitted model with (C) or without (D) prior incidence beyond the village. Data points in red fall outside the 95% prediction interval (PI).

**Figure S8:**
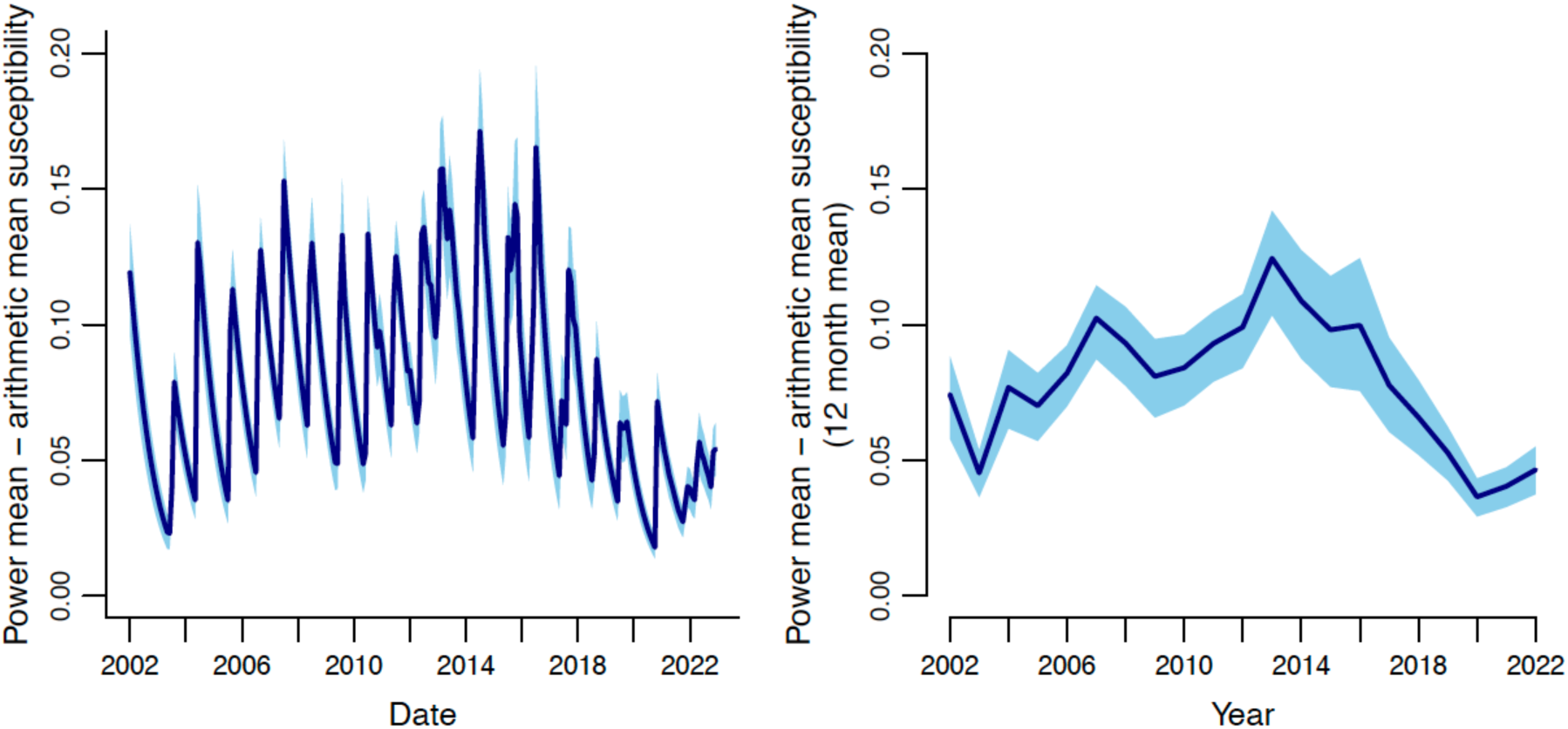
Difference between power mean and arithmetic mean susceptibility. Difference between power mean susceptibility calculated over all villages in the district using fitted values of *p* from the model without prior incidence beyond the focal village (Fig. 4C) minus the arithmetic mean for each month (A) or averaged over each year (B).

**Figure S9:**
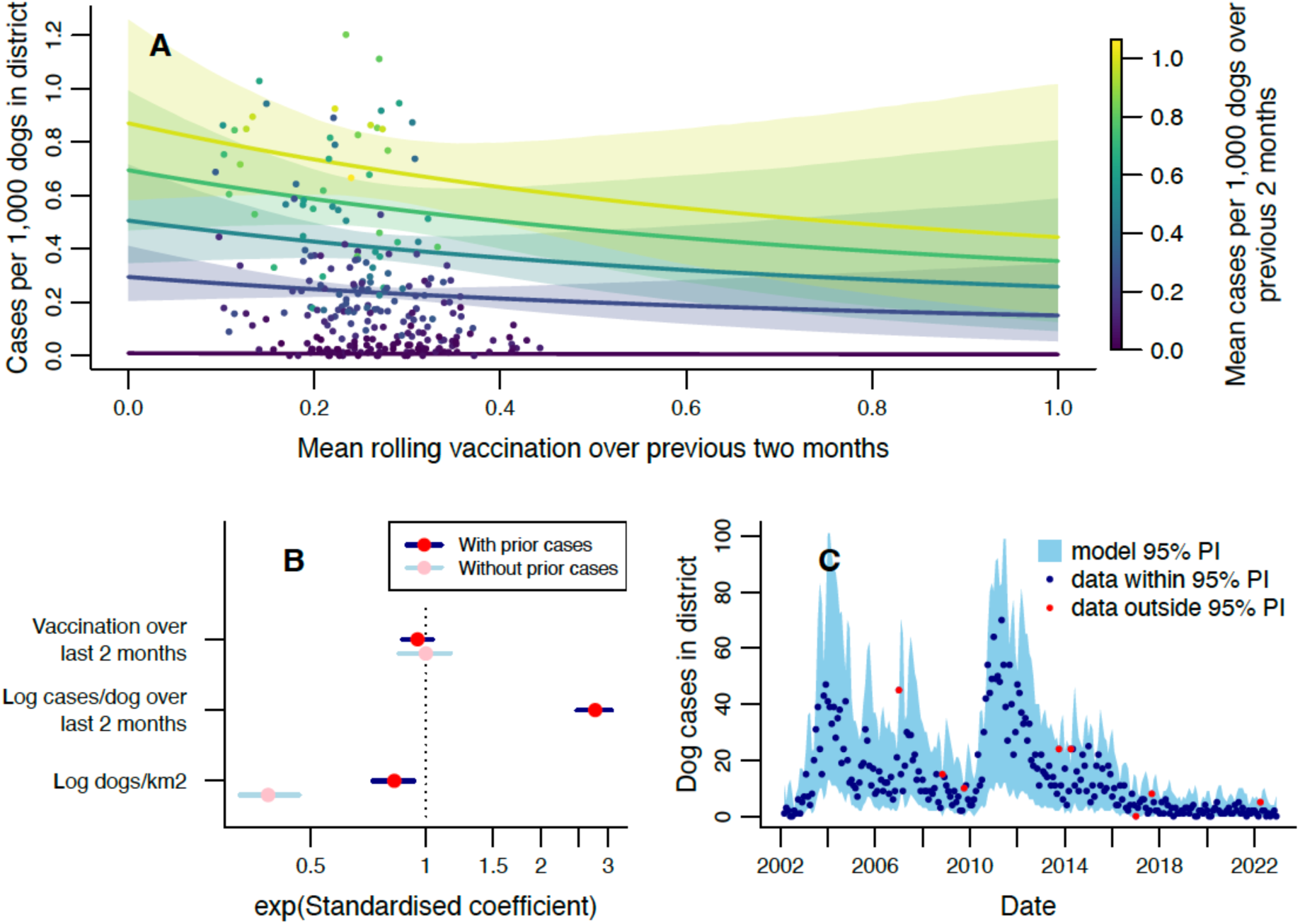
Modelling monthly dog rabies cases in Serengeti District at district level. A) Expected cases/dog (number of dog cases normalised by dog population) in the district this month for different mean rolling vaccination coverages and mean cases/dog in the prior 2 months. Shaded areas show 95% CrIs, points show the data, and predictions were obtained using the average value of dog density. B) Exponentiated standardised values of the coefficients estimated for each explanatory variable, with 95% CrIs. Coefficients obtained for a version of the model fitted without prior cases/dog as an explanatory variable are included for comparison. See Table S2 for tabulated parameter values. C) Comparison of observed monthly dog cases (points) with the 95% prediction interval from the fitted model. Data points in red fall outside the 95% prediction interval (PI).

**Figure S10:**
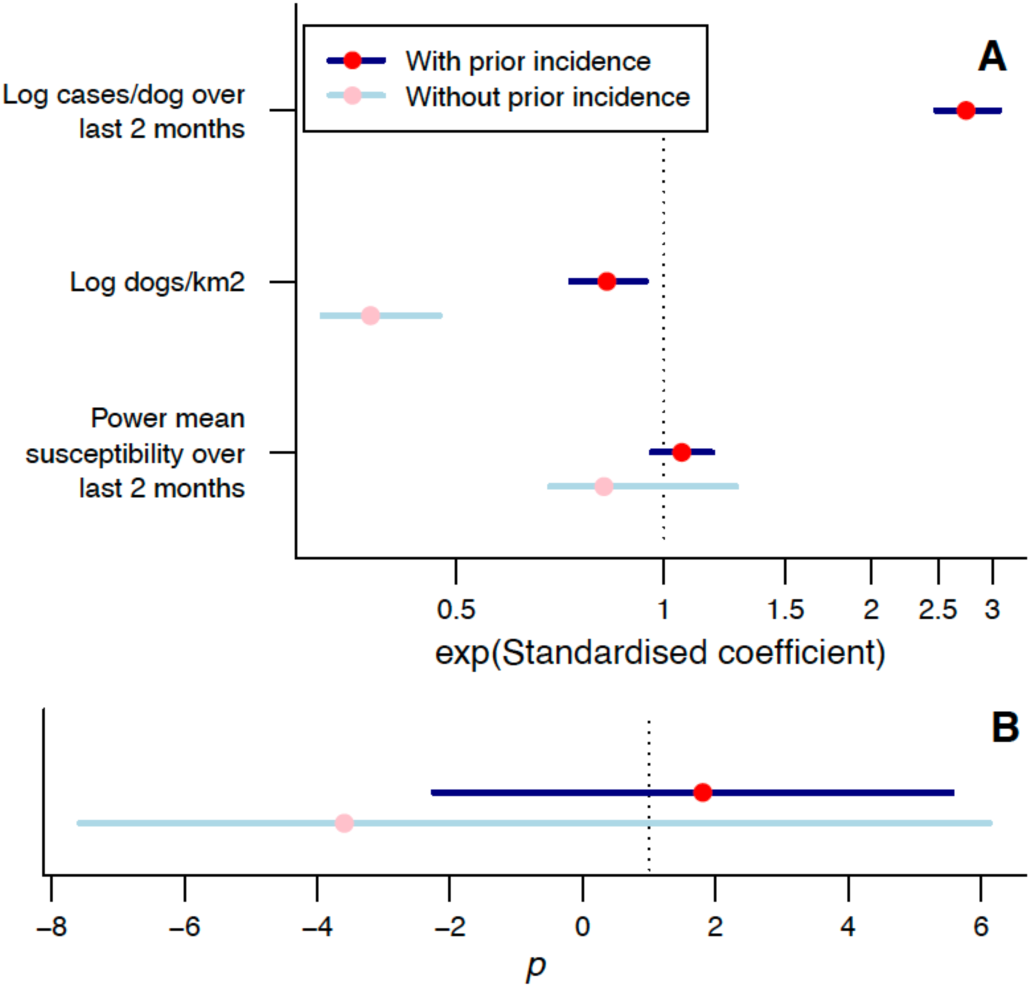
Using power mean susceptibility to model impacts of heterogeneity in rolling vaccination on district-level incidence. A) Exponentiated standardised estimated coefficients for each explanatory variable, with 95% CrIs. B) Estimated power *p* used to calculate power mean susceptibility. In A-B, estimates from models with and without effects of prior incidence are shown. See Table S2 for tabulated parameter values.

**Figure S11:**
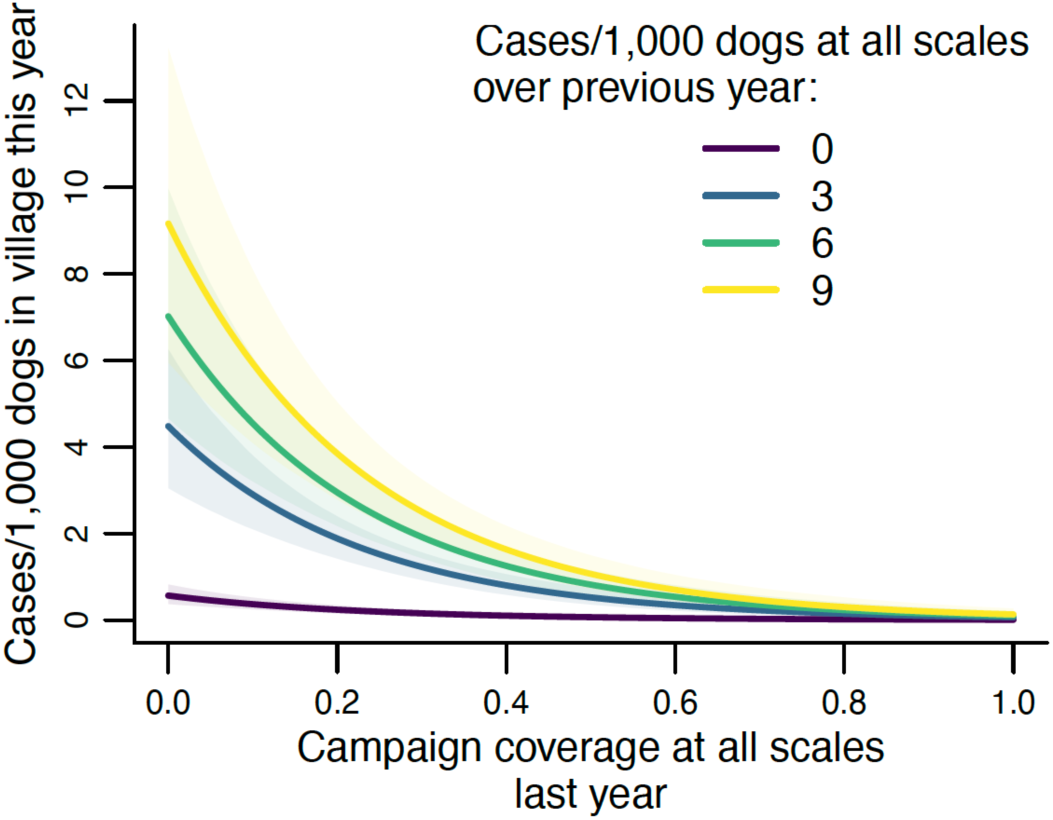
Annual village-level GLMM for current cases per dog. Lines show the expected cases per 1,000 dogs (number of dog cases normalised by dog population) in a village this year for different campaign vaccination coverages and cases per 1,000 dogs in the previous year. Prior incidence values were chosen to represent the range observed at district level. Shaded areas show 95% CrIs, and predictions were obtained using average values of unspecified explanatory variables. See Table S3 for tabulated parameter values.

**Figure S12:**
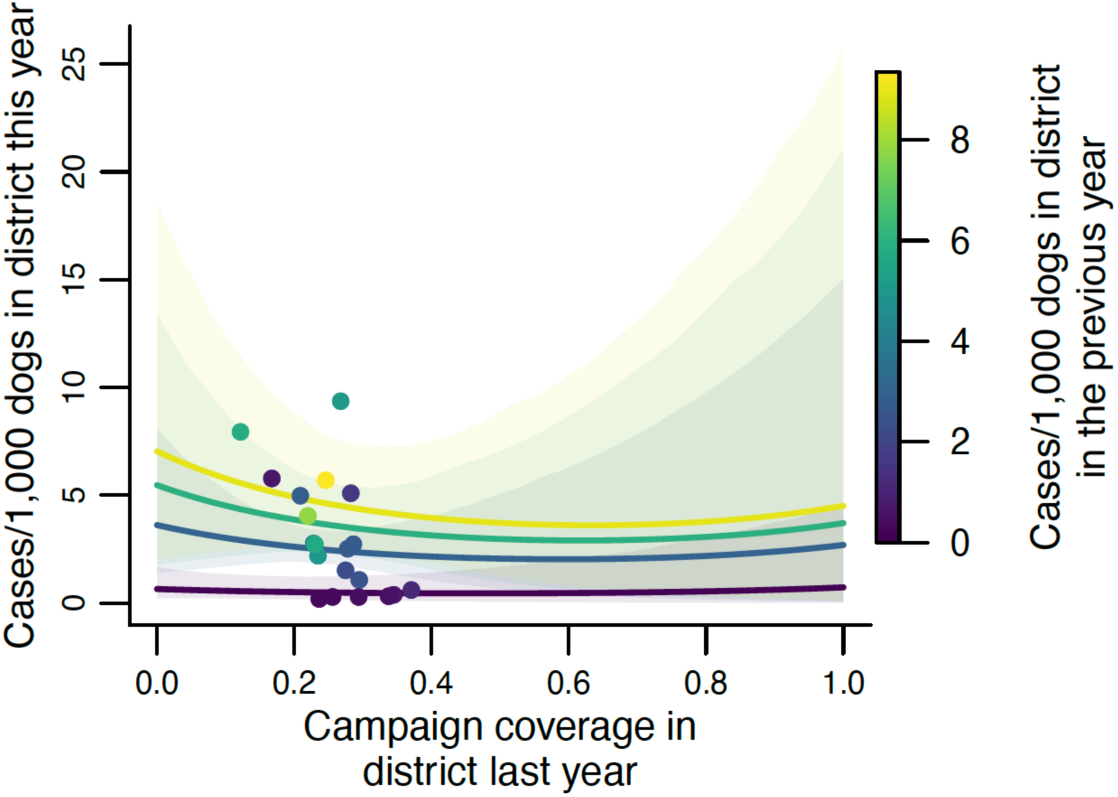
Annual district-level GLM for cases per dog. Lines show the expected cases per 1,000 dogs (number of dog cases normalised by dog population) in the district this year for different campaign vaccination coverages and cases per 1,000 dogs in the previous year. Prior incidence values were chosen to represent the range observed at district level. Shaded areas show 95% CrIs, and predictions were obtained using average values of unspecified explanatory variables. See Table S4 for tabulated parameter values.

**Figure S13:**
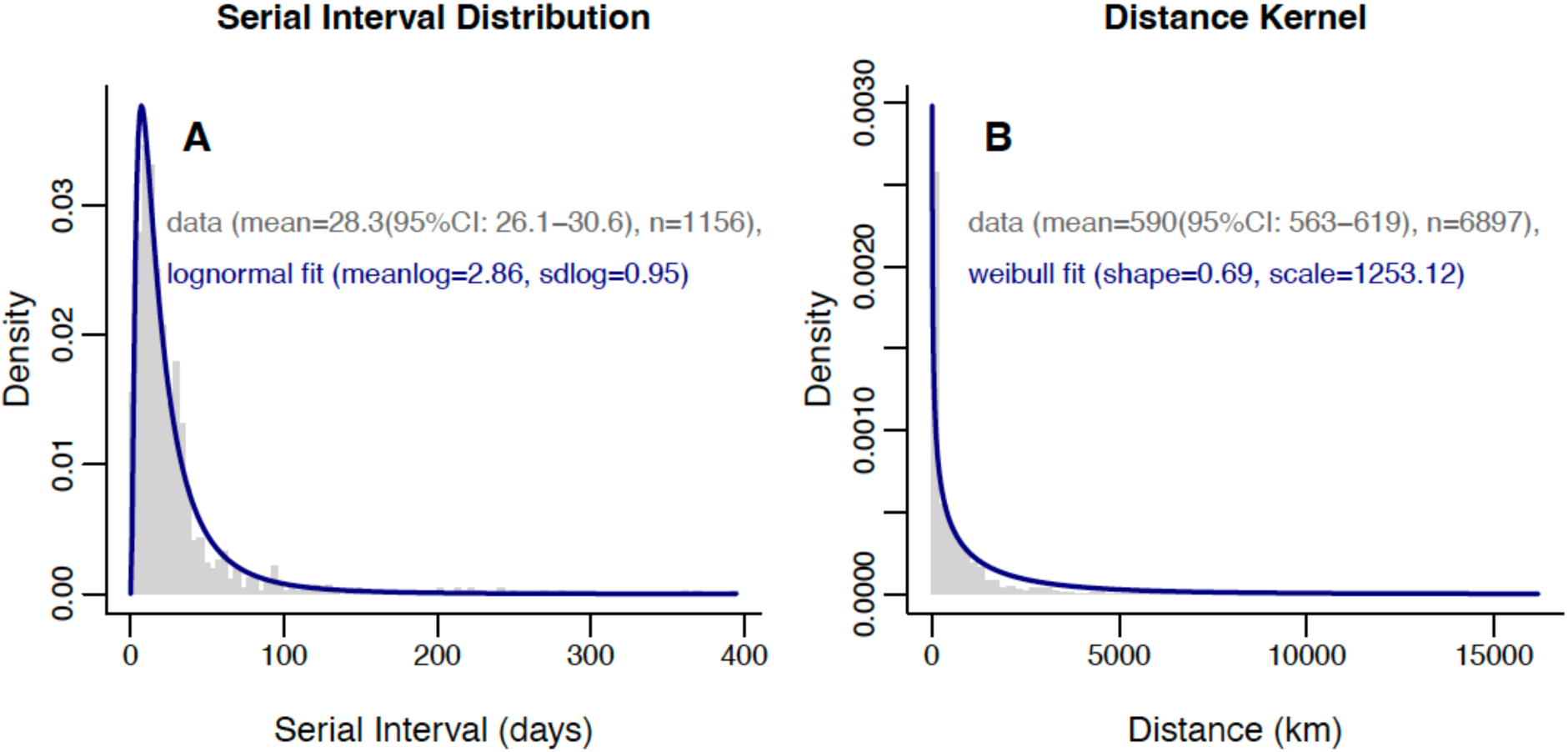
Serial interval distribution and distance kernel. A) Histogram of serial intervals calculated from contact tracing data (grey bars), with fitted lognormal distribution (blue line). B) Histogram of distances between the starting location of a case and the locations of its contacts (grey bars), with fitted weibull distribution (blue line).

**Figure S14:**
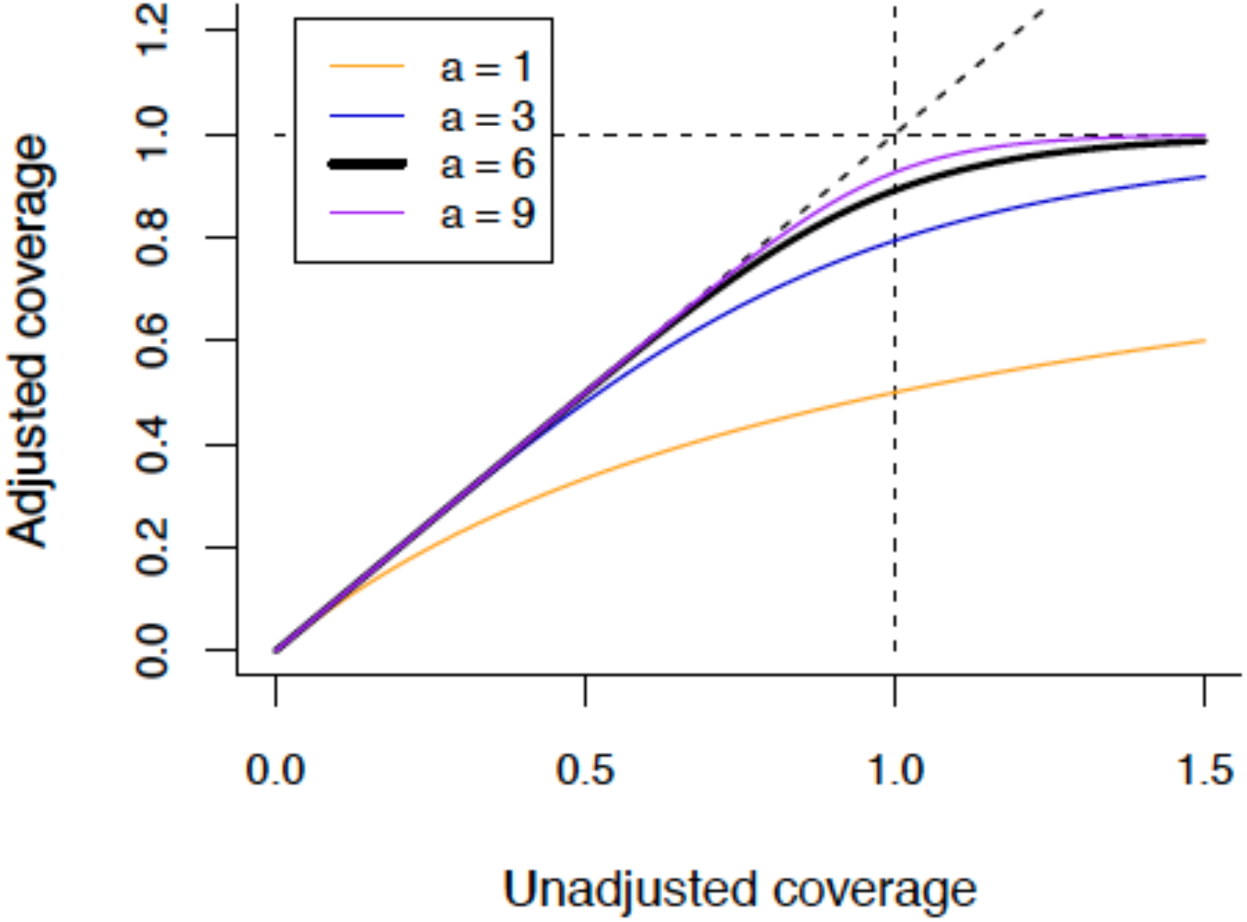
Bounding the proportion of dogs vaccinated between zero and one. Illustration of the function *b*(*x*)*=x/*((1+*x^a^*)^1/*a*^) used to bound coverage estimates below one. Throughout our analyses, we set a=6 when applying this function, but the impact of using alternative values is shown here.

### Supplementary Tables

**Table S1:**
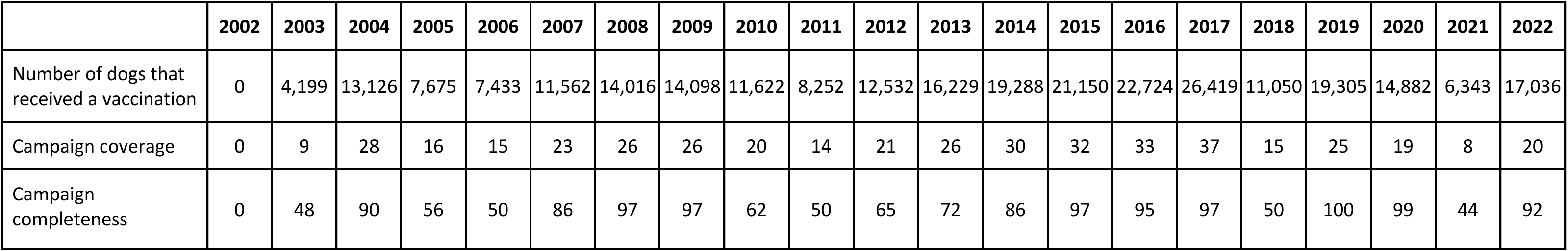
Vaccination of Serengeti District by year. Numbers of dogs that received a vaccination in each year, campaign coverage (percentage of the district dog population vaccinated in campaigns each year), and campaign completeness (percentage of villages in the district that held a campaign in each year).

**Table S2:** Parameter estimates for monthly district-level GLMs for cases/dog in the district in the current month. 95% credible intervals (CrIs) in brackets. Coefficients for fixed effects where the 95% CrI does not include zero are marked *. Predictions from model 1 (and coefficients for model 2) are presented in Fig. S9). Parameters from models 3 and 4 are illustrated in Fig. S10. Vaccination, susceptibility and cases/dog variables are all averages over the prior two months.

| Parameter | 1. Full model without power mean | 2. Without power mean or prior cases/dog | 3. Full model with power mean | 4. Power mean without prior cases/dog |
| --- | --- | --- | --- | --- |
| Intercept | 1.52 (0, 3.1) | 6.67 (4.03, 9.31) | 0.5 (-2.15, 2.99) | 8.01 (0.17, 11.42) |
| Rolling vaccination coverage | -0.75 (-2.06, 0.55) | -0.06 (-2.18, 2.13) |  |  |
| Susceptibility |  |  | 0.92 (-0.5, 2.46) | -1.32 (-2.98, 4.19) |
| Log cases/dog | 0.78 (0.7, 0.86)* |  | 0.78 (0.7, 0.86)* |  |
| Log dogs/km<sup>2</sup> | -0.98 (-1.59, -0.38)* | -4.71 (-5.59, -3.81)* | -0.94 (-1.54, -0.35)* | -4.89 (-5.72, -3.68)* |
| size (negative binomial distribution parameter) | 4.76 (3.55, 6.32) | 1.3 (1.06, 1.57) | 4.79 (3.58, 6.31) | 1.36 (1.11, 1.65) |
| $p$ (power used in calculating power means) | | | 2.02 (-1.74, 5.73) | -3.06 (-7.32, 6.32) |

**Table S3:**
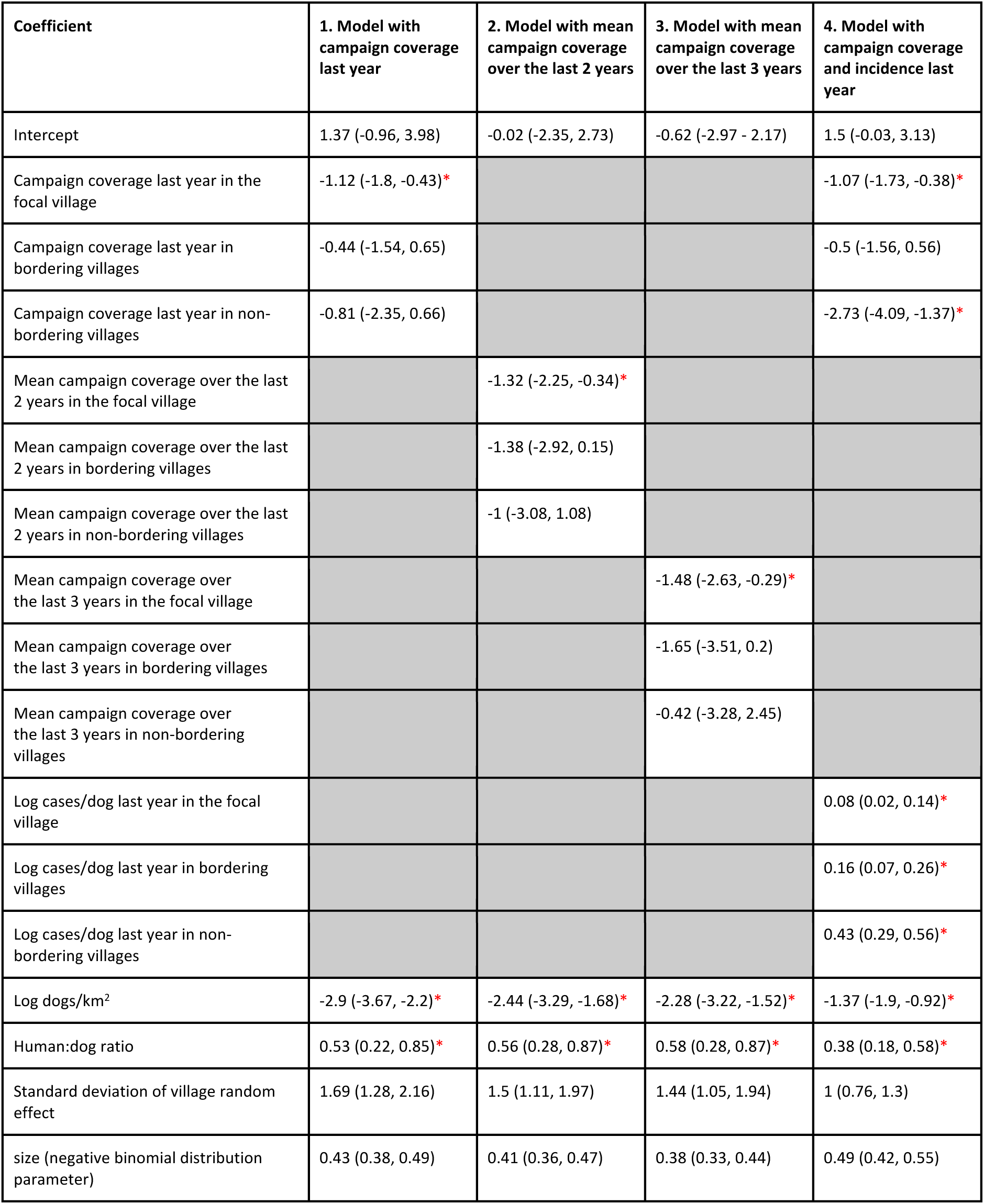
Coefficients for the annual village-level negative binomial GLMMs. 95% credible intervals in brackets. Coefficients for fixed effects where the 95% CrI does not include zero are marked *. Predictions from the model 4 (including campaign coverage and cases/dog in the last year as explanatory variables) are presented in Fig. S11.

| Coefficient | 1. Model with campaign coverage last year | 2. Model with mean campaign coverage over the last 2 years | 3. Model with mean campaign coverage over the last 3 years | 4. Model with campaign coverage and incidence last year |
| --- | --- | --- | --- | --- |
| Intercept | 1.37 (-0.96, 3.98) | -0.02 (-2.35, 2.73) | -0.62 (-2.97 - 2.17) | 1.5 (-0.03, 3.13) |
| Campaign coverage last year in the focal village | -1.12 (-1.8, -0.43)* |  |  | -1.07 (-1.73, -0.38)* |
| Campaign coverage last year in bordering villages | -0.44 (-1.54, 0.65) |  |  | -0.5 (-1.56, 0.56) |
| Campaign coverage last year in non-bordering villages | -0.81 (-2.35, 0.66) |  |  | -2.73 (-4.09, -1.37)* |
| Mean campaign coverage over the last 2 years in the focal village |  | -1.32 (-2.25, -0.34)* |  |  |
| Mean campaign coverage over the last 2 years in bordering villages |  | -1.38 (-2.92, 0.15) |  |  |
| Mean campaign coverage over the last 2 years in non-bordering villages |  | -1 (-3.08, 1.08) |  |  |
| Mean campaign coverage over the last 3 years in the focal village |  |  | -1.48 (-2.63, -0.29)* |  |
| Mean campaign coverage over the last 3 years in bordering villages |  |  | -1.65 (-3.51, 0.2) |  |
| Mean campaign coverage over the last 3 years in non-bordering villages |  |  | -0.42 (-3.28, 2.45) |  |
| Log cases/dog last year in the focal village |  |  |  | 0.08 (0.02, 0.14)* |
| Log cases/dog last year in bordering villages |  |  |  | 0.16 (0.07, 0.26)* |
| Log cases/dog last year in non-bordering villages |  |  |  | 0.43 (0.29, 0.56)* |
| Log dogs/km<sup>2</sup> | -2.9 (-3.67, -2.2)* | -2.44 (-3.29, -1.68)* | -2.28 (-3.22, -1.52)* | -1.37 (-1.9, -0.92)* |
| Human:dog ratio | 0.53 (0.22, 0.85)* | 0.56 (0.28, 0.87)* | 0.58 (0.28, 0.87)* | 0.38 (0.18, 0.58)* |
| Standard deviation of village random effect | 1.69 (1.28, 2.16) | 1.5 (1.11, 1.97) | 1.44 (1.05, 1.94) | 1 (0.76, 1.3) |
| size (negative binomial distribution parameter) | 0.43 (0.38, 0.49) | 0.41 (0.36, 0.47) | 0.38 (0.33, 0.44) | 0.49 (0.42, 0.55) |

**Table S4:** Coefficients for the annual district-level negative binomial GLMs. 95% credible intervals in brackets. Coefficients for fixed effects where the 95% CrI does not include zero are marked *. Predictions from model 4 (including campaign coverage and cases/dog in the last year as explanatory variables) are presented in Fig. S12.

| Coefficient | 1. Model with campaign coverage last year | 2. Model with mean campaign coverage over the last 2 years | 3. Model with mean campaign coverage over the last 3 years | 4. Model with campaign coverage and incidence last year |
| --- | --- | --- | --- | --- |
| Intercept | 12.29 (4.84, 19.74) | 13.8 (5.6, 22.2) | 14.64 (5.61, 23.25) | 9.51 (4, 15.09) |
| Campaign coverage last year | 0.55 (-4.1, 4.93) |  |  | -1.08 (-4.66, 2.42) |
| Mean campaign coverage over the last 2 years |  | 0.6 (-5.75, 6.87) |  |  |
| Mean campaign coverage over the last 3 years |  |  | 1.92 (-6.96, 10.67) |  |
| Log cases/dog last year |  |  |  | 0.55 (0.21, 0.89)* |
| Log dogs/km<sup>2</sup> | -5.7 (-8.04, -3.29)* | -6.15 (-8.79, -3.45)* | -6.5 (-9.16, -3.6)* | -3.72 (-5.84, -1.64)* |
| size (negative binomial distribution parameter) | 2.08 (1.03, 3.6) | 2.08 (0.99, 3.59) | 2.03 (0.94, 3.56) | 3.25 (1.5, 5.81) |

**Table S5:** Fitted distributions for incubation period (units=days), serial interval (units=days) and distance kernel (units=metres). For each epidemiological variable calculated from the contact tracing data with sample size n, we fitted gamma, lognormal and Weibull distributions. Estimates of the parameters for each distribution are provided. The best fitting model for each epidemiological variable based on AIC is highlighted in bold. The 95th, 97.5th and 99th percentiles of each distribution are given.

|  | n | Distribution | Parameter 1:<br>Name | Parameter 1:<br>Value (95%CI) | Parameter 2:<br>Name | Parameter 2:<br>Value (95%CI) | AIC | 95th<br>percentile | 97.5th<br>percentile | 99th<br>percentile |
| --- | --- | --- | --- | --- | --- | --- | --- | --- | --- | --- |
| **Incubation**<br>**Period** | 1,212 | Gamma | shape | 1.18 (1.1, 1.26) | rate | 0.04 (0.04, 0.05) | 10421 | 77.2 | 93.9 | 115.9 |
|  |  | **Lognormal** | **meanlog** | **2.82 (2.77, 2.87)** | **sdlog** | **0.95 (0.91, 0.99)** | **10159** | **80.1** | **108.0** | **152.9** |
|  |  | Weibull | shape | 1 (0.96, 1.05) | scale | 27.27 (25.78, 28.89) | 10440 | 81.6 | 100.5 | 125.4 |
| **Serial**<br>**Interval** | 1,156 | Gamma | shape | 1.18 (1.1, 1.27) | rate | 0.04 (0.04, 0.05) | 10022 | 79.9 | 97.2 | 119.9 |
|  |  | **Lognormal** | **meanlog** | **2.86 (2.81, 2.92)** | **sdlog** | **0.95 (0.91, 0.99)** | **9781** | **83.4** | **112.5** | **159.3** |
|  |  | Weibull | shape | 1 (0.96, 1.05) | scale | 28.29 (26.73, 30.09) | 10041 | 84.5 | 104.0 | 129.8 |
| **Distance**<br>**kernel** | 6,897 | Gamma | shape | 0.58 (0.55, 0.6) | rate | 0.00038 (0.00035, 0.00041) | 45177 | 5596.7 | 7216.4 | 9407.9 |
|  |  | Lognormal | meanlog | 6.45 (6.4, 6.49) | sdlog | 1.74 (1.69, 1.8) | 45321 | 11129.8 | 19284.6 | 36540.9 |
|  |  | **Weibull** | **shape** | **0.69 (0.67, 0.71)** | **scale** | **1253.12 (1192.2, 1315.93)** | **45143** | **6142.8** | **8304.8** | **11453.2** |

